# Miniscope Zero: a fully wireless, single-cell-resolution miniature microscope for imaging neural dynamics in freely behaving animals

**DOI:** 10.64898/2026.09.05.745822

**Authors:** Takuya Sasatani, Marcel Brosch, Zhe Dong, Jonny Saunders, Pingping Zhao, Megha Sehgal, Lukas T. Oesch, Federico Sangiuliano Jimka, Hemal Semwal, Aparajeeta Guha, Hamid Chorsi, Karina Keus, Blake A. Madruga, Alcino J. Silva, Anne K. Churchland, Peyman Golshani, Daniel Aharoni

## Abstract

Imaging neural fluorescence dynamics during unconstrained behavior remains a major challenge in neuroscience. Head-mounted miniature one-photon microscopes enable in vivo recordings from freely behaving animals. Still, their power and data requirements typically necessitate tethers or heavy batteries, which constrain experiments and may introduce behavioral artifacts. We present Miniscope Zero, a fully wireless miniature microscope platform combining quasistatic cavity resonance (QSCR) wireless power transfer with a high-bandwidth optical data link. The system delivers >500 mW of receiver power across a 2,500 cm² behavioral arena and streams imaging data at 8 Mbps, supporting real-time data acquisition. Miniscope Zero provides a typical field of view of 600 µm × 600 µm and a 2.6-fold higher light-collection efficiency than the UCLA Miniscope v4. We demonstrate wireless CA1 GCaMP6f recordings during open-field navigation, enclosed-maze exploration, simultaneous multi-animal imaging, and extended three-dimensional behavior, enabling long-duration cellular-resolution imaging without tether-induced constraints.

---

Understanding how neural population dynamics support complex behavior requires tools that record cellular activity during freely moving behavior. Genetically encoded calcium indicators enable large-scale monitoring of neuronal populations over extended periods^1^, but conventional benchtop microscopes generally require head fixation. Head-mounted miniature microscopes bring cellular-resolution imaging to freely behaving animals^2,3^, and open-source platforms, including the UCLA Miniscope^4^, FinchScope^5^, mini-mScope^6^, Kiloscope/Featherscope^7^, and NiNscope^8^, have enabled studies of neural activity during navigation^9,10^, learning^11^, and memory^12^.

Extended recordings during complex behavior nevertheless remain constrained by miniature imaging hardware. Most systems remain tethered, using custom cable assemblies or coaxial cables^8,9^ for power and data transmission, or optical fibers^7,13,14^ for relaying light to and from off-scope hardware. These tethers introduce translational drag, rotational torque, and cable-management constraints that can bias behavior^15–17^. These limitations are especially notable in enclosed structures, multi-animal experiments, and three-dimensional environments, where cables can restrict movement or become entangled. Several untethered miniscopes eliminate the physical tether but remain dependent on finite onboard batteries, with data either stored locally or transmitted wirelessly. However, battery-powered systems trade recording duration against head-mounted mass^5,9^; onboard storage prevents real-time visualization and adjustment of recording parameters^9,18^; and previous wireless data implementations have operated at reduced image resolution and frame rate^19^. Fully wireless systems providing both wireless power and data transmission have been developed for applications with considerably lower power and bandwidth requirements, including optogenetic stimulation^20,21^ and fiber photometry^22,23^. Miniature calcium imaging, however, imposes substantially greater simultaneous power and data-throughput demands, precluding incremental scaling of these architectures^24^. Fully wireless miniature microscopy therefore requires the development and integration of wide-area power delivery, high-throughput data transmission and compact head-mounted optics.

Here we present Miniscope Zero—named for its zero-tether operation—a fully wireless one-photon calcium-imaging platform for recordings in freely behaving animals. Miniscope Zero combines wide-area wireless power based on quasistatic cavity resonance (QSCR)^25,26^ with a low-power, high-bandwidth optical communication link. The standard head-mounted configuration weighs 4.12 g, although this value does not capture the elimination of tether-induced drag and torque, whose effects cannot be represented by a single equivalent mass^17^. The QSCR transmitter delivers >500 mW of receiver power across a 2,500 cm² behavioral arena, providing sufficient margin for continuous microscope operation without a large onboard battery. An 8-Mbps optical uplink supports real-time imaging at 200 × 200 pixels and 20 frames per second, while an infrared downlink enables adjustment of sensor gain, excitation intensity, and imaging region of interest. The wireless power and wavelength-separated optical data architecture supports simultaneous recording from multiple animals. A redesigned optical path provides a typical 600 μm × 600 μm field of view and collects 2.6 times as much emitted light as our previous-generation wired miniscope, the UCLA Miniscope v4. By eliminating both power and high-bandwidth data tethers, Miniscope Zero enables cellular-resolution neural imaging during increasingly unconstrained behavior.

## Results

Miniscope Zero integrates the head-mounted microscope, QSCR power interface, optical data I/O, and acquisition software into an end-to-end platform (Fig. 1a–d). An onboard battery buffers fluctuations in current draw and transient reductions in wireless power coupling. Initial recordings used a 50-mAh battery (1 g), whereas subsequent testing showed that an 11-mAh battery (0.33 g) still supports continuous operation in the standard 4.12-g configuration. We validate the system by imaging GCaMP6f-expressing CA1 neurons during open-field navigation and exploration of enclosed mazes with vertical and overlapping paths, obtaining stable wireless recordings with robust spatial tuning (Fig. 1e,f). We further compare behavioral effects with a conventional tethered Miniscope v4 (Fig. 4e–g) and demonstrate extended operation in multi-animal and complex environments (Fig. 1g,h). Representative Miniscope Zero recordings across the experimental environments are shown in Supplementary Video 1.

**Figure 1.**
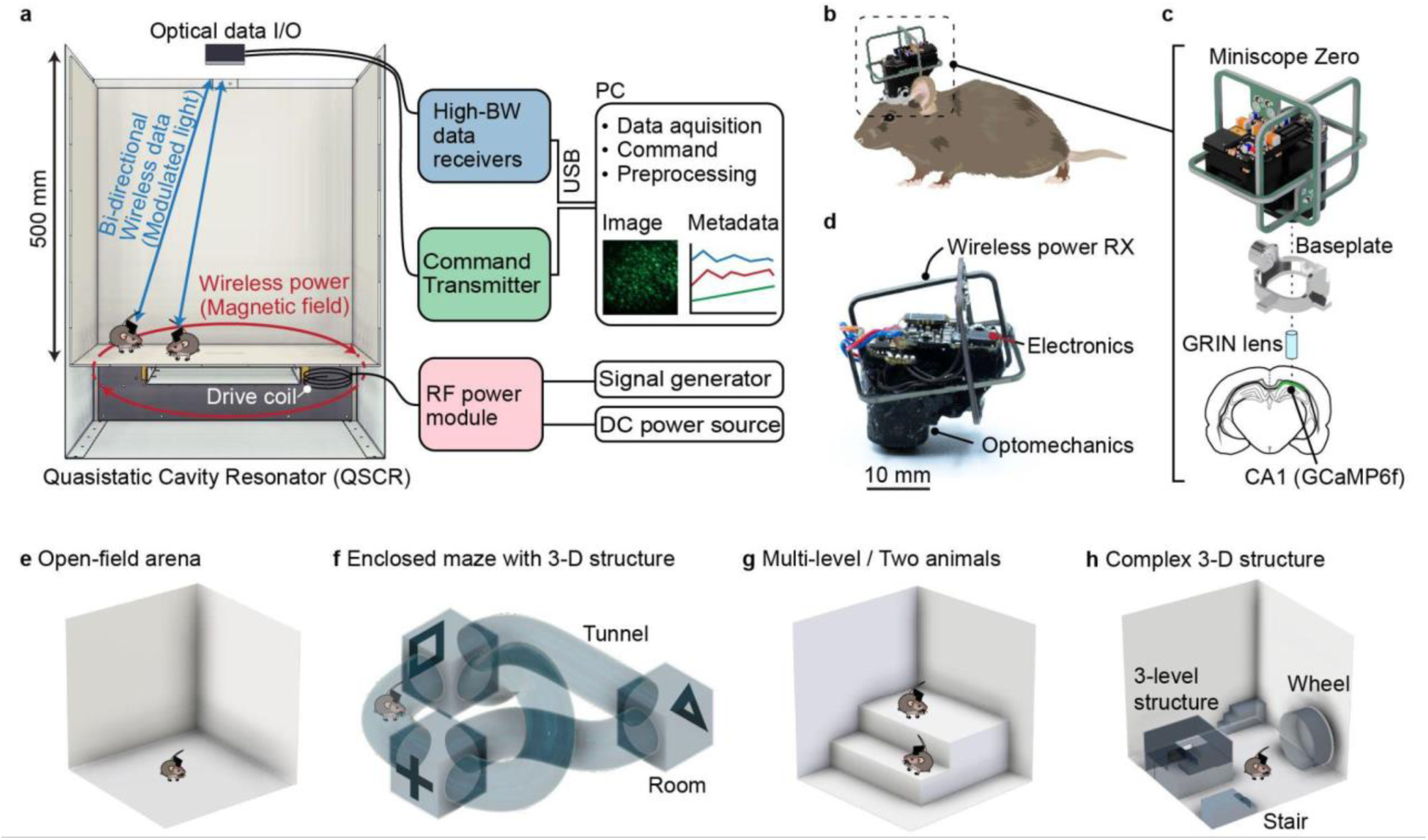
Miniscope Zero wireless neural-imaging platform and experimental configurations. **a,** System overview. A QSCR cavity provides wide-area wireless power, while bidirectional optical communication supports imaging-data transmission and remote device control. **b,** Freely behaving mouse carrying Miniscope Zero. **c,** Exploded view showing the head-mounted device, baseplate, implanted gradient-index (GRIN) lens and GCaMP6f-expressing neurons in dorsal CA1. **d,** Assembled Miniscope Zero, integrating the wireless-power receiver, imaging and communication electronics, and optomechanical components. **e–h,** Experimental configurations: **e,** open-field arena; **f,** enclosed maze with overlapping tunnels and rooms; **g,** multilevel arena for simultaneous two-animal imaging; and **h,** complex environment containing a three-level structure, stairs, and a running wheel. The configurations in **f–h** illustrate settings in which conventional tethers restrict enclosed and multi-animal behavior, whereas fully wireless operation with Miniscope Zero enables imaging throughout these environments. Mouse and brain illustrations adapted from SciDraw under CC BY 4.0; full attribution is provided in the Acknowledgments.

### High-efficiency optical design

The Miniscope Zero uses a redesigned emission path that improves fluorescence collection relative to the UCLA Miniscope v4 while preserving single-cell resolution (Fig. 2 and Extended Data Fig. 1). The collection numerical aperture (NA) is 0.45 (versus 0.3 for the Miniscope v4), and the magnification is 1.6× (versus 3×), which together increase collection efficiency and concentrate emitted photons onto fewer sensor pixels. The design is intended to achieve a comparable signal at lower excitation power. Under matched acquisition conditions using a microLED panel to isolate the emission path from excitation, mean 8-bit pixel intensity was 38.3 ± 4.3 for Miniscope v4 and 98.2 ± 7.7 for Miniscope Zero (mean ± s.d.), corresponding to a 2.6-fold increase in per-pixel signal (Fig. 2e).

**Figure 2.**
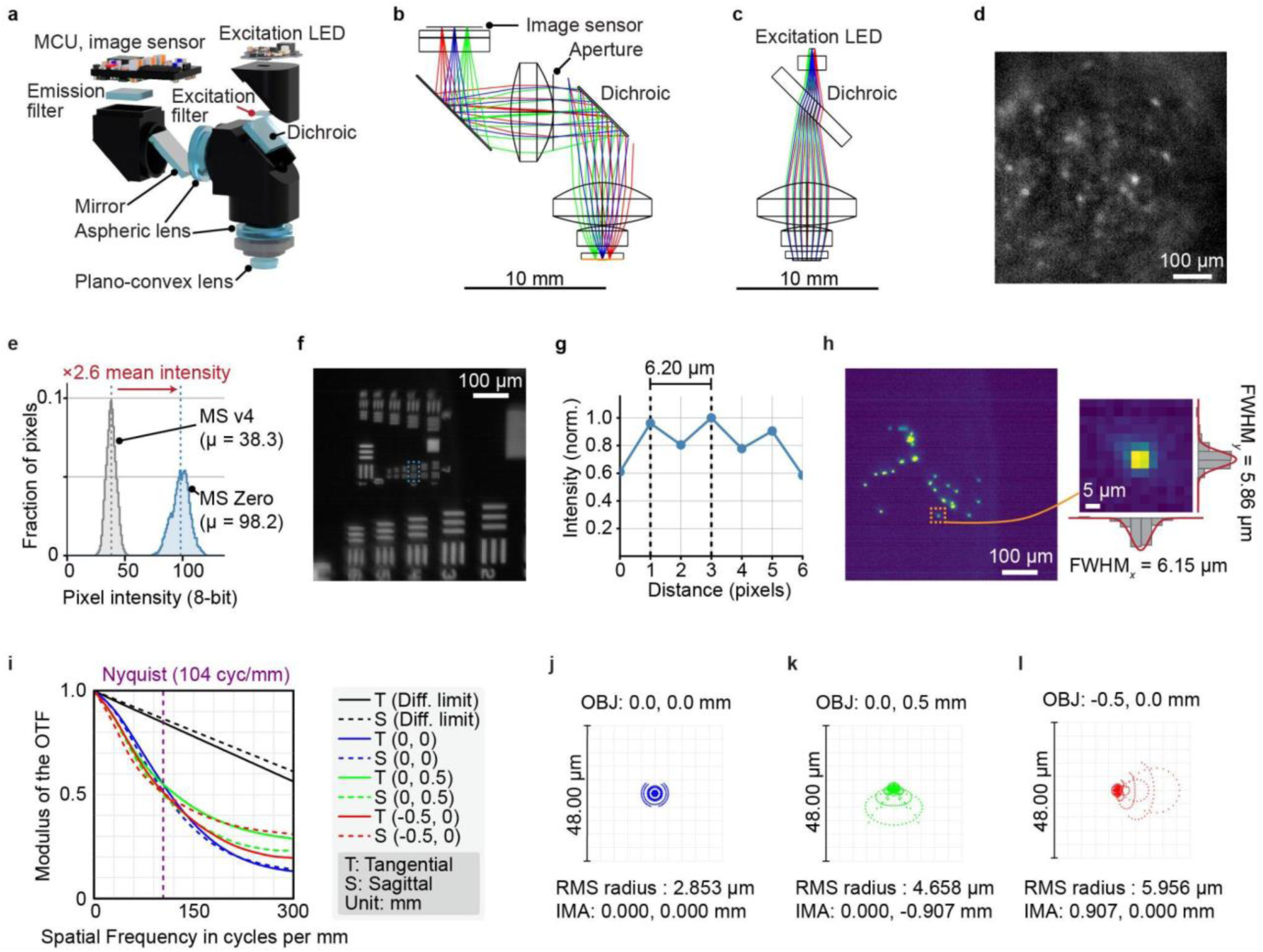
Optical design and characterization of Miniscope Zero. **a,** Exploded view of the Miniscope Zero optical assembly. The emission path is folded laterally by the dichroic and planar mirror, enabling a high numerical aperture while maintaining a low-profile geometry. **b,c,** Ray-tracing models of the emission (**b**) and excitation (**c**) optical paths. **d,** Example frame from the preprocessed calcium-imaging video showing GCaMP6f-expressing CA1 neurons recorded with Miniscope Zero. Preprocessing was performed as described in the Methods. **e,** Emission-collection comparison between Miniscope Zero and the UCLA Miniscope v4 optical paths under matched acquisition conditions using an emission-only microLED target. Distributions show 8-bit pixel intensities, with Miniscope Zero producing a 2.6-fold higher mean intensity than Miniscope v4. **f,g,** Spatial-resolution characterization using a negative USAF 1951 resolution target. Miniscope Zero resolved Group 7, Element 3, corresponding to a 3.1-μm line width and 161.3 lp mm⁻¹, in both horizontal and vertical orientations. The corresponding line-pair period, 6.2 µm, is reported as the resolution limit. **f,** Representative target image. **g,** Normalized intensity profile across the resolved line pattern. **h,** Measured lateral point-spread function using 200 nm fluorescent microspheres. Left, field of view containing multiple beads; right, magnified view of a single bead with horizontal and vertical intensity profiles. **i,** Simulated tangential and sagittal modulation transfer functions at the field center and at two positions 0.5 mm from the optical axis. The purple dashed line indicates the image-sensor Nyquist frequency, and the black curves indicate the diffraction limit. **j–l,** Simulated geometric spot diagrams at the field center (**j**) and at two positions 0.5 mm from the optical axis (**k,l**).

The increased collection efficiency is achieved while decreasing overall device height. The optical path is folded twice by 90° using a dichroic mirror and a planar mirror, distributing the optics laterally rather than vertically, resulting in a 12 mm height, lower than the 22 mm height of the Miniscope v4, and creating a low center of mass. Miniscope Zero also resolves structures smaller than a neuronal soma. On a USAF 1951 target, it resolved a 3.1 µm line width, corresponding to a 6.2 µm line-pair period, which we report as the resolution limit (Fig. 2f,g). In agreement, the lateral point-spread function measured from fluorescent microspheres had a full width at half maximum (FWHM) of 6.05 ± 0.54 µm (mean ± s.d., *n* = 10; Fig. 2h). These values are below the typical diameter of a neuronal soma (10–20 µm) and well below the effective neuronal footprint after scattering in tissue.

### Wide-range wireless power transfer

A central challenge for fully wireless miniature microscopy is delivering the few hundred milliwatts required for complementary metal–oxide–semiconductor (CMOS) imaging and LED excitation while keeping the head-mounted device within a few grams. Conventional inductive links^27,28^ typically deliver high power only over limited areas, as in charging pads, making continuous power delivery during behavior difficult^29^. Miniscope Zero addresses this challenge using QSCR^25,26,30^, in which the behavioral arena forms part of a resonant structure that generates a magnetic field throughout the experimental volume, reducing sensitivity to the animal’s position (Fig. 3a,b and Extended Data Fig. 2). Three orthogonal head-mounted receiver coils further reduce orientation sensitivity as the animal turns, rears and moves through the environment (Fig. 1c,d and Extended Data Fig. 3d). The system therefore functions as an environmental power source for the microscope rather than a point-to-point charger.

**Figure 3.**
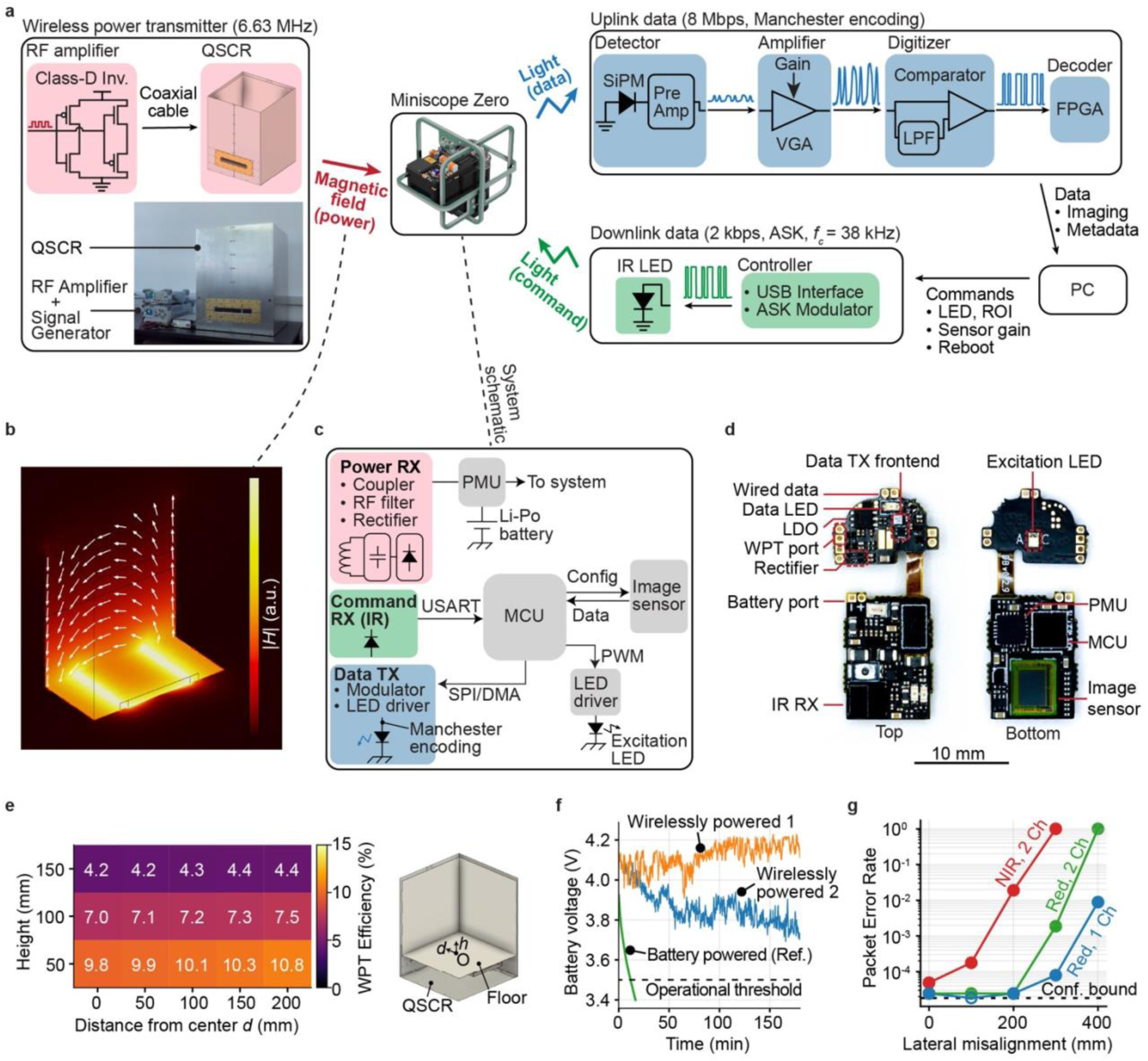
Architecture and performance of the Miniscope Zero wireless platform. **a,** System architecture integrating wireless power transfer at 6.63 MHz using a QSCR, an 8-Mbps Manchester-encoded optical uplink for imaging data and metadata, and a 2-kbps infrared downlink for device control. **b,** Simulated magnetic-field distribution within the QSCR at the operating frequency. **c,** Miniscope Zero electronics architecture. **d,** Top and bottom views of the Miniscope Zero circuit-board assembly, showing the principal power, imaging, and communication components. **e,** Measured wireless power-transfer efficiency as a function of receiver height and lateral distance from the arena center. Transfer efficiency varied weakly with lateral position at each height and decreased as the receiver was elevated above the arena floor. **f,** Battery-voltage trajectories during simultaneous wireless operation of two Miniscope Zero test devices for 3 h, compared with a battery-only reference device. The dashed line indicates the 3.5 V operational threshold used during experiments. **g,** Packet error rate as a function of lateral misalignment between the optical transmitter and receiver for single-channel and wavelength-separated two-channel configurations. For trials with no observed packet errors, the dashed line indicates the approximate 95% upper confidence bound, 3/*N*.

The QSCR transmitter operates at 6.63 MHz and provides wireless power across a 2,500 cm² arena. Following established practice for characterizing wireless-power links, we evaluated RF power-transfer efficiency under the receiver-load condition that maximizes transfer efficiency^25,26,31^. Efficiency was approximately 10% at a typical mouse head height of 50 mm and 4% at 150 mm above the floor, indicating usable coupling across the vertical range relevant to freely moving mice (Fig. 3e). Circuit-model analysis attributed 90.6–96.0% of transfer losses to the drive coil and QSCR resonator rather than the head-mounted receiver coil. In separate measurements under representative operating conditions, the system delivered more than 500 mW to the receiver, providing substantial margin over the approximately 200 mW required to operate Miniscope Zero. Each of the three head-mounted receiver coils weighs approximately 200 mg, and a small onboard battery buffers transient reductions in coupling caused by animal motion or orientation.

### High-bandwidth optical communication link

We developed a miniaturized optical link to avoid the power and footprint costs associated with previous high-speed wireless communication implementations for neural imaging (Fig. 3a,c,d). An 8-Mbps uplink streams 8-bit images at 200 × 200 pixels and 20 FPS, together with metadata for frame reconstruction and device monitoring. A 2-kbps infrared downlink enables adjustment of sensor gain, excitation intensity, and imaging region of interest during acquisition.

For single-animal recordings, multiple receivers can capture the same optical stream, and packet metadata enable post hoc combination of the receiver streams to extend coverage beyond that of a single receiver (Extended Data Fig. 4b,e). Receiver modules can therefore be distributed across larger or geometrically complex arenas. For simultaneous multi-animal recordings, wavelength-separated red and near-infrared uplinks, together with matched optical filters, provide independent data channels.

At a packet error rate threshold of 10^−3^, corresponding to an expected frame loss rate of approximately 0.8% for frames comprising eight packets, a single receiver provided a lateral coverage diameter of approximately 70 cm, equivalent to an area of approximately 3,800 cm², in the single-channel configuration (Fig. 3g). In the two-channel configuration, the corresponding diameter was approximately 30 cm, equivalent to approximately 700 cm². The reduced coverage in the two-channel configuration reflects the angular sensitivity of the wavelength-selective filters and can be extended by adding receiver modules.

### Wireless imaging of CA1 place cells during freely behaving open-field navigation

Hippocampal place cells can encode an animal’s spatial location and are thought to contribute to cognitive maps supporting navigation and memory. To evaluate Miniscope Zero during freely behaving experiments, we imaged GCaMP6f-expressing dorsal CA1 pyramidal neurons while mice explored an open-field arena.

#### Place cell recording

We recorded CA1 activity while mice explored a 500 × 500 mm open field (Fig. 4a–d). Miniscope Zero was mounted above a chronically implanted relay GRIN lens using a baseplate (Fig. 1c), and post hoc histology confirmed that the lens was positioned over GCaMP6f-expressing tissue in CA1 (Fig. 4d).

**Figure 4.**
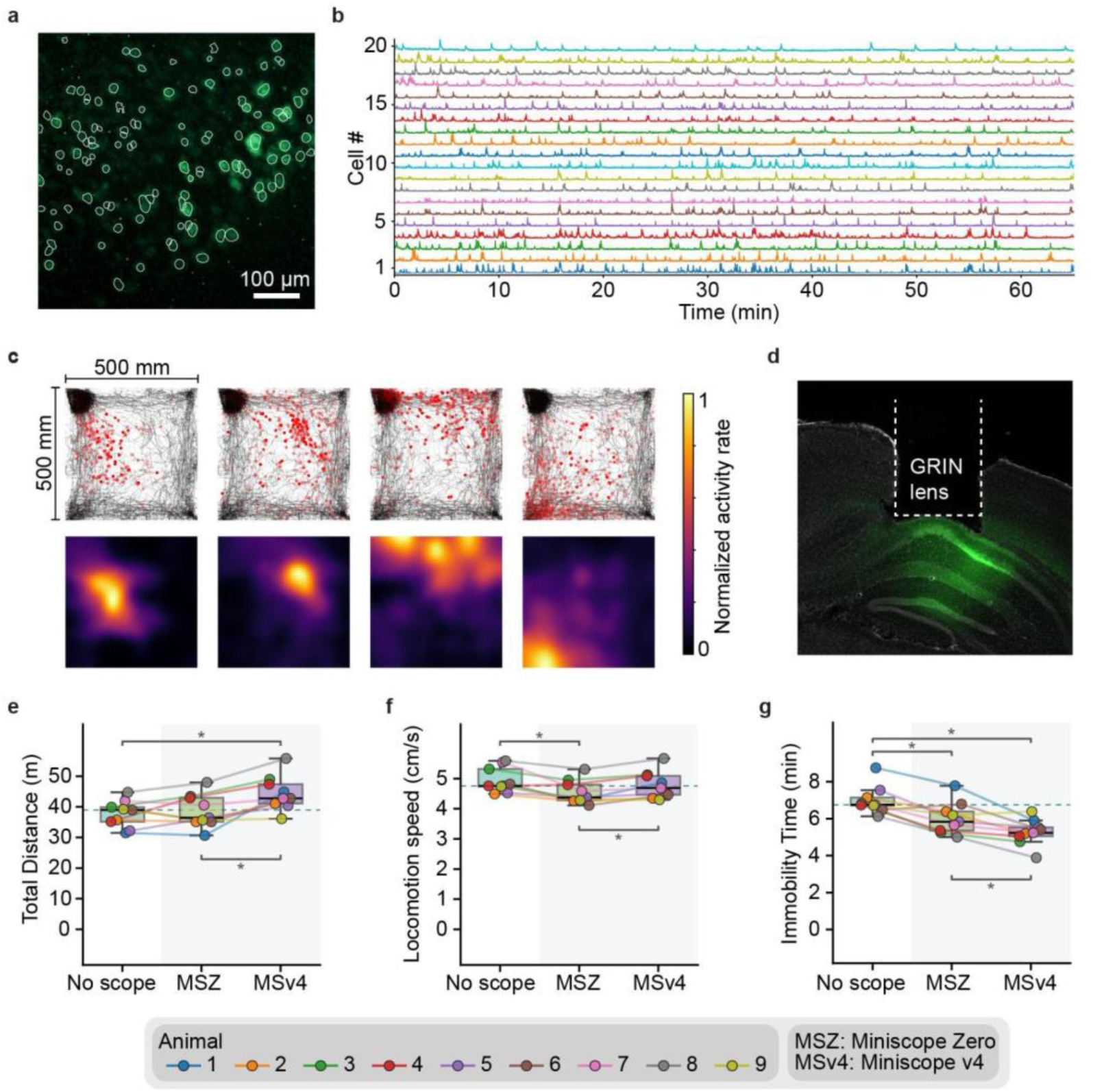
Wireless CA1 calcium imaging and behavioral comparison during open-field navigation. **a,** Maximum-intensity projection from an example 60-min open-field recording, with spatial footprints of neurons identified using MiniAn outlined. In this session, 134 neurons were detected, of which 75 were classified as place cells. **b,** Activity traces from 20 simultaneously recorded CA1 neurons. **c,** Spatial activity of four example CA1 neurons during open-field exploration. Top, animal trajectories with calcium-event locations overlaid; bottom, corresponding occupancy-normalized activity-rate maps. **d,** Histological section showing GCaMP6f expression in dorsal CA1 beneath the implanted gradient-index (GRIN) lens. **e–g,** Within-animal behavioral comparison under three conditions: no-miniscope control (no scope), Miniscope Zero (MSZ), and Miniscope v4 (MSv4). Nine mice were recorded in a 300 mm × 300 mm open-field arena. **e,** Total distance traveled. **f,** Locomotion speed, calculated from frames in which the animal was mobile. **g,** Total immobility time. Each point represents the median value for one mouse in each condition (*n* = 9), and lines connect measurements from the same animal. Boxes show the median and interquartile range across mice (whiskers 1.5 × IQR). The dashed horizontal line marks the no-scope median and the shaded band groups the two scope conditions. Conditions were compared using Friedman tests, followed by two-sided Wilcoxon signed-rank tests with Holm–Bonferroni correction when the omnibus test was significant. Significant pairwise comparisons are indicated above the plots; \**P* < 0.05.

In the open-field experiments, we recorded three mice in five 30-min sessions each, and one of these mice in an additional 60-min session, yielding 16 sessions in total. We treated each session as an independent dataset for quantification^32^. In the example 60-min session, we identified 134 neurons using MiniAn^33^, of which 75 met the shuffle-based spatial-information and stability criteria for place cells. Combining deconvolved activity with tracked position revealed spatially localized activity maps, with distinct neurons preferentially active in different regions of the arena (Fig. 4a–c). Across the 16 sessions, mice visited 79.9 ± 9.9% of the 1 × 1 cm spatial bins. We detected 135 ± 55 neurons per session (mean ± s.d.; 2,156 neurons in total), of which 36.7 ± 13.3% were classified as place cells.

#### Miniscope behavior comparison

We next tested whether the wireless configuration altered behavior relative to a conventional tethered miniscope. We recorded 9 mice in a 300 mm × 300 mm open-field arena under three within-animal conditions: no-scope control, Miniscope Zero, and Miniscope v4 (Fig. 4e–g). We compared distance traveled, locomotion speed, and immobility. Miniscope Zero weighed 4.12 g, including the microscope body, wireless power receiver coils, and an 11-mAh battery, whereas Miniscope v4 weighed 2.8 g without its tether.

Both devices altered behavior, but along different axes. Mice carrying a Miniscope v4 traveled approximately 10% farther than controls (median, 42.8 versus 39.0 m; Friedman *P* = 0.004, Wilcoxon/Holm *P* = 0.023) and approximately 17% farther than mice carrying a Miniscope Zero (36.4 m; *P* = 0.012). Distance traveled with Miniscope Zero was indistinguishable from that of the control (−7%; *P* = 0.73). Miniscope Zero reduced locomotion speed by 8% relative to control (4.4 versus 4.8 cm s⁻¹; Friedman *P* = 0.004; Wilcoxon/Holm *P* = 0.016) and was approximately 7% slower than Miniscope v4 (4.69 cm s⁻¹; *P* = 0.012), whereas Miniscope v4 was indistinguishable from control (−2%; *P* = 0.20). Both scopes reduced immobility: by approximately 23% with Miniscope v4 (5.2 versus 6.8 min; Friedman *P* < 0.001, Wilcoxon/Holm *P* = 0.012) and by approximately 14% with Miniscope Zero (5.8 min; *P* = 0.016), with Miniscope v4 approximately 10% below Miniscope Zero (*P* = 0.016). Each device altered two of the three metrics. These findings suggest that tether drag, torque, or other mechanical factors can contribute to behavioral disruption beyond the effects of head-mounted mass alone.

### Wireless imaging in tether-incompatible environments

To test Miniscope Zero in configurations in which a tethered miniscope would interfere directly with the behavioral environment or the animal’s movement, we evaluated wireless imaging in three environments (Figs. 5 and 6): an enclosed structured maze, a complex environment containing multiple three-dimensional structures, and a shared multilevel arena containing two simultaneously imaged mice.

**Figure 5.**
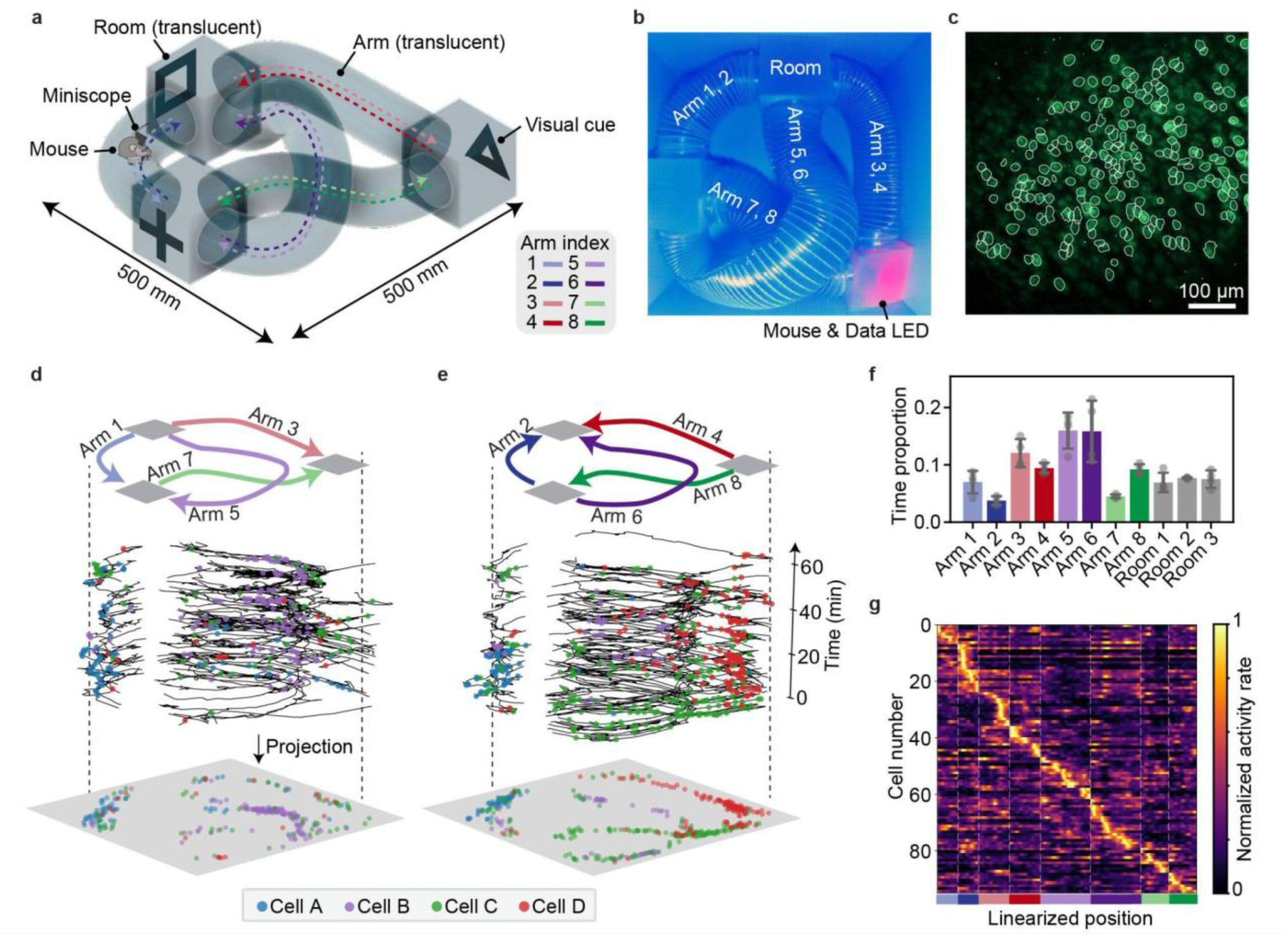
Wireless CA1 calcium imaging during navigation in an enclosed structured maze. a,b, Schematic (**a**) and top view (**b**) of the maze, comprising translucent rooms connected by enclosed, overlapping arms. Conventional tethers would restrict movement through these paths. Colored dashed lines indicate the eight directed arms used for linearized-position analysis. **c,** Maximum-intensity projection from an example 60-min recording of GCaMP6f-expressing CA1 neurons, with MiniAn-identified neuronal footprints outlined. In this session, 218 neurons were detected, of which 95 were classified as place cells. **d,e,** Spatial activity of four example CA1 neurons during directed-arm traversals. Top, analyzed arm trajectories; middle, animal position over time, with colored points indicating calcium events from the example neurons; bottom, projection of event locations onto the maze layout. **f,** Proportion of recording time spent in each maze arm and room. Bars show mean ± s.d.; gray points indicate individual sessions; *n* = 4 sessions from 2 mice. **g,** Occupancy-normalized activity-rate maps for the 95 place cells identified in the example session, plotted along the linearized maze position. Each row represents a neuron, normalized to its peak activity and ordered by peak position; the colored bar indicates the arm identity. Mouse illustrations adapted from SciDraw under CC BY 4.0; full attribution is provided in the Acknowledgments.

**Figure 6.**
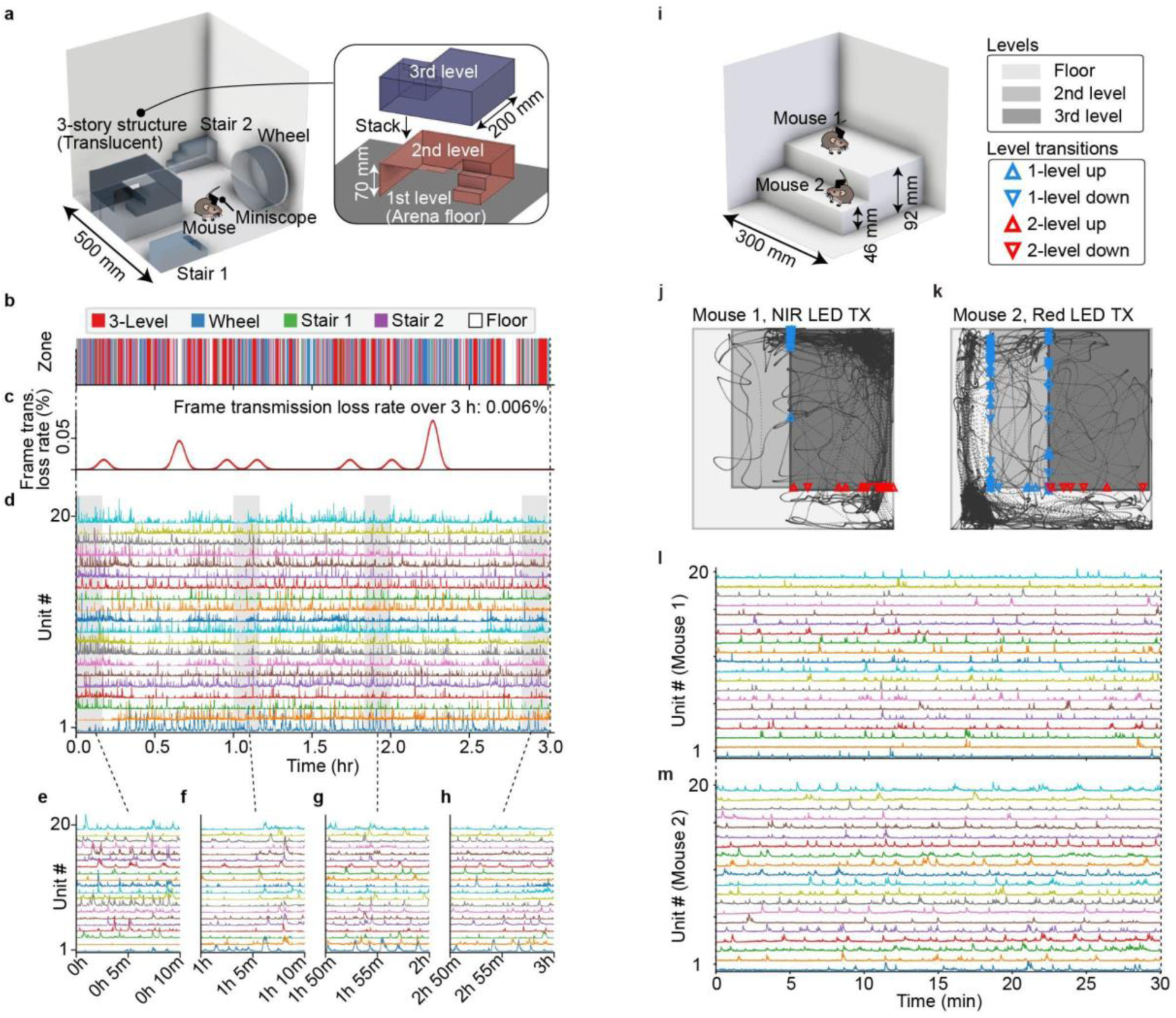
Demonstrations of wireless imaging capabilities in tether-incompatible environments. **a–h,** 3-h single-animal imaging in a complex environment. **a,** Schematic of the environment used for the 3-h recording, comprising a translucent three-level structure, two stairways, a running wheel, and open floor space. **b,** Behavioral-zone assignment throughout the recording, indicating occupancy of the three-level structure, wheel, stairways, and floor. **c,** Frame transmission loss rate during the same session, calculated in 0.5-min bins and smoothed with a Gaussian kernel with a 5-min full width at half maximum. Total frame transmission loss over the full 3 h recording was 0.006%. **d,** Calcium activity traces from 20 representative CA1 neurons across the full 3-h recording. Shaded regions indicate the intervals expanded in **e–h**. **e–h,** Selected 10-min excerpts showing calcium transients at higher temporal resolution at different times during the recording. **i–m,** Simultaneous two-animal imaging in a multilevel arena. **i,** Schematic of the arena. **j,k,** LED-tracked trajectories of mouse 1 using the near-infrared optical uplink (**j**) and mouse 2 using the red optical uplink (**k**) during a 30-min recording. Blue markers indicate transitions between adjacent levels, red markers indicate direct transitions between non-adjacent levels, and marker orientation indicates upward or downward movement. The two mice completed 38 and 75 inter-level transitions, respectively. **l,m,** Simultaneously acquired calcium activity traces from 20 representative CA1 neurons in mouse 1 (**l**) and mouse 2 (**m**). The experiments in **a–m** were performed as qualitative demonstrations of the device’s capabilities during prolonged exploration and simultaneous multi-animal behavior. Mouse illustrations adapted from SciDraw under CC BY 4.0; full attribution is provided in the Acknowledgments.

#### Recording in an enclosed maze

We first recorded dorsal CA1 activity while mice navigated a 500 × 500 mm structured maze of rooms connected by an enclosed tunnel system (Fig. 5). The translucent maze walls visually enclosed the tunnels while diffusely transmitting the data-LED signal, allowing optical data reception as the mouse moved through enclosed and vertically overlapping paths that would make tethered operation impractical. Across four recording sessions from two mice, data acquisition continued for the full intended duration despite repeated transitions between enclosed tunnels and rooms. Across the first 30 min of each recording, combined frame transmission loss averaged 0.3% (Extended Data Fig. 5a).

The recordings remained suitable for downstream analysis and yielded 125 ± 77 detected neurons per session (mean ± s.d.; 500 neurons in total). Applying spatial-information and stability criteria, 38.8 ± 8.7% of cells were classified as place cells. In the example 60-min session, 218 neurons were detected, of which 95 met the place-cell criteria. The mouse traveled throughout the maze, and place fields spanned the linearized trajectory of the tunnel system (Fig. 5c–g). These results demonstrate that Miniscope Zero can maintain physiologically interpretable neural recordings as animals navigate enclosed and vertically overlapping paths.

#### Continuous 3-h imaging in a complex environment

We next tested whether wireless operation could be maintained during extended exploration of an open environment containing a three-level structure with platforms 70 and 140 mm above the arena floor, two 70-mm-high stairways, and a running wheel (Fig. 6a). Wireless power was maintained and CA1 activity was recorded throughout the 3-h session as the mouse repeatedly moved among these behavioral zones (Fig. 6b). Despite changes in head height, orientation and optical line of sight, dual-receiver reconstruction limited combined frame transmission loss to 0.006% over the full session, and the reconstructed images supported extraction of calcium events (Fig. 6c–h).

#### Simultaneous two-animal imaging in a multilevel arena

We next tested whether two Miniscope Zero devices could be powered and recorded simultaneously while both animals moved freely through a shared multilevel arena (Fig. 6i). Wavelength-separated red and near-infrared optical uplinks enabled separate recovery of the two imaging streams. During the 30-min session, both mice repeatedly moved between platforms positioned 46 and 92 mm above the arena floor, including direct transitions between non-adjacent levels (Fig. 6j,k). Both imaging streams supported extraction of CA1 activity traces throughout the session (Fig. 6l,m). These results demonstrate simultaneous wireless power delivery and cellular-resolution imaging from two freely moving animals.

### Wireless operation reliability

We characterized wireless-operation reliability across multiple environments and in both single- and dual-animal configurations. We transmitted battery and rectified-input voltages within the optical metadata stream, enabling real-time monitoring of device power state and manual adjustment of QSCR transmitter input. This telemetry-guided control maintained operation during recordings lasting 30 min to 3 h in open-field, enclosed-maze, complex-environment and multilevel configurations (Figs. 4–6), a 3-h two-animal wireless-power test (Fig. 3f) and a 9-h single-animal in vivo capability test (Extended Data Fig. 6).

For single-animal recordings in 500 × 500 mm arenas, we combined dual-receiver streams so a frame was classified as lost only when neither receiver received all eight packets. This metric reports packet-level delivery; frames that arrived complete but were subsequently rejected by image quality control are accounted for separately (Methods). Across the first 30 min of each recording, combined frame transmission loss was 0.002% in the open arena, 0.003% in the complex environment, and 0.3% in the enclosed maze (Extended Data Fig. 5a), despite transient occlusions, posture changes, and variation in optical line of sight.

For simultaneous two-animal recordings in 300 × 300 mm arenas, wavelength-separated red and near-infrared uplinks were acquired as independent streams, each on its own dedicated receiver. Across the first 30 min, all frames were delivered complete in the red channel in both the open-field and multilevel arena, whereas the near-infrared channel showed 0.05% and 0.6% incomplete frames, respectively (Extended Data Fig. 5b). In the multilevel arena, both mice repeatedly climbed between platforms 4.6 and 9.2 cm above the floor, testing operation across changes in receiver height and orientation.

During the 9-h in vivo device capability test, dual-receiver combining reduced frame transmission loss from approximately 3.4% and 0.05% in the individual receiver streams to 0.005% in the combined stream. Together, these results show that QSCR power delivery and optical receiver redundancy support long-duration wireless operation in single- and multi-animal experiments.

## Discussion

Miniscope Zero provides, to our knowledge, the first miniature calcium-imaging platform to combine wide-range wireless power with real-time high-bandwidth wireless data transmission, addressing two constraints that commonly limit long-duration and multi-animal recordings: dependence on an external power tether or finite head-mounted battery, and dependence on a high-bandwidth imaging-data tether or onboard-only data storage. By using QSCR-based wireless power, a small rechargeable onboard buffer, and an optical data link, the system supports real-time cellular-resolution imaging without making recording duration primarily dependent on head-mounted battery capacity or onboard storage. This architecture enabled stable CA1 population recordings during both open-field navigation and enclosed-maze exploration. Paired behavioral comparisons showed that although Miniscope Zero was heavier, it did not produce a uniformly larger behavioral disruption: each device altered two of three locomotor metrics. Head-mounted mass is therefore not the sole determinant of behavioral disruption; tether drag and torque likely contribute independently.

The present implementation is optimized for experiments within a QSCR cavity, but the architecture is modular. Because the wireless-power system primarily replaces a large battery or power tether, related designs could be adapted to other head-mounted or body-mounted devices with compatible power requirements. Calcium imaging is relatively power- and data-bandwidth-demanding compared with many other sensing and stimulation modalities, suggesting that the Miniscope Zero’s wireless architecture could be adapted to a wide range of miniaturized tools. Future work could also explore transmitter geometries beyond the cavity form factor used here, including designs tailored to larger spaces, home-cage recordings, or specialized behavioral apparatuses. More broadly, Miniscope Zero provides a platform for adapting power delivery, data transmission, and optical imaging to the behavioral environment, enabling cellular-resolution studies of brain–behavior relationships under increasingly unconstrained conditions.

## Methods

Data acquisition, frame reconstruction, and offline combination of receiver streams were implemented in our MIO (Miniscope I/O) Python package. Spatial coding analyses were implemented in our CaMAP Python package. Repository links are provided in the Code availability section.

### Animal preparation

All experimental protocols were approved by the Chancellor’s Animal Research Committee at the University of California, Los Angeles (protocol number 2006-066), and were carried out in accordance with U.S. National Institutes of Health guidelines for the care and use of laboratory animals.

This study used two cohorts of adult C57BL/6J mice. The imaging cohort consisted of three male mice (12 weeks old) used to record Ca²⁺ activity from dorsal CA1 (dCA1) pyramidal cells. After implantation, animals for imaging were single-housed to protect the implant. A second cohort of ten mice (5 male, 5 female; 12 weeks old) was used for a behavioral comparison between the UCLA Miniscope v4 and Miniscope Zero; these animals received only a baseplate implantation (no craniotomy, viral injection, or GRIN lens) and were not used for calcium imaging. Animals for purely behavioral studies were group-housed. All mice were housed on a 12:12 h light/dark cycle with food and water available ad libitum.

### Surgical implantation for hippocampal imaging

Mice underwent three survival surgeries before behavioral training. During the first surgery, animals were anesthetized with isoflurane (5% for induction, 1–2% for maintenance), and a small incision was made above the dorsal CA1. 500 nL of AAV-Syn-GCaMP6f (AAV1.Syn.GCaMP6f.WPRE.SV40, Addgene #100837, titer 1.1 × 10^13^ vg mL^−1^) was injected unilaterally at the following coordinates relative to bregma: AP −2.1 mm, ML 2.0 mm, DV −1.65 mm, via a glass capillary (Drummond, 3-000-203-G/X) mounted on a Nanoject III (Drummond Scientific) at a rate of 60 nL min^−1^. The injection needle was left in place for 10 min after injection to allow for diffusion before slow withdrawal. The scalp was closed using Vetbond (3M, 1468SB).

Three to five days after viral injection, mice were again anesthetized for GRIN lens implantation^34^. One stainless steel screw (00-96 × 1/16, Plastics One) was inserted into the skull to anchor the implant. A 0.75 cm diameter circle of skin was removed to expose the skull, and the cortical tissue overlying the hippocampus was aspirated using a blunted 27- and then 30-gauge needle until the alveus was exposed. After confirming hemostasis, a 1.0-mm-diameter GRIN lens (Inscopix, 1050-004595) was lowered to the following coordinates relative to bregma: AP −2.1 mm, ML 1.8 mm, DV −1.25 mm. The lens was cemented in place with cyanoacrylate glue (Krazy Glue) and dental cement (Ortho-Jet), along with the dorsal surface of the skull and skull screw, to ensure implant stability. The lens was protected with Kwik-Sil (World Precision Instrument).

Four to five weeks after viral injection, once GCaMP6f expression had stabilized, a milled aluminum baseplate was cemented above the lens to ensure stable miniscope mounting. The miniscope was mounted on the stereotaxic arm and positioned above the implanted lens at the focal plane where Ca²⁺ fluorescence and identifiable cell bodies were in focus. The baseplate was then cemented in place with dental cement, the miniscope was removed, and the cement was allowed to cure. A protective cap was fixed to the baseplate between recording sessions. After recovery from baseplating, mice began behavioral training.

### Histology

At the end of the experiment, mice were euthanized with a lethal dose of isoflurane, followed by an intraperitoneal injection of 1 mL of pentobarbital, and then transcardially perfused with 20 mL of 1× phosphate-buffered saline (PBS) followed by 20 mL of 4% paraformaldehyde in 1× PBS to fix the brain tissue. Brains were dehydrated in 30% sucrose solution and sectioned at 60 μm thickness on a cryostat (Leica), mounted on slides, and imaged on a confocal microscope (Zeiss) to confirm green fluorescent protein (GFP) expression and GRIN lens placement.

### Miniscope Zero design

The head-mounted microscope integrates an epifluorescence optical path, a CMOS image sensor, an excitation LED, a low-power microcontroller (Microchip, ATSAMD51J20A-UUT-EFP), a wireless power receiver, power management circuitry, and bidirectional optical communication modules into a compact head-mounted assembly. The microcontroller configures the image sensor, controls frame acquisition, regulates excitation timing, and coordinates data transmission. The device also retains a wired diagnostic interface to support testing, troubleshooting, and data acquisition during stereotaxic baseplating, manual focus adjustment, or other fixed preparations.

The optical assembly was designed to increase emission collection while maintaining a compact head-mounted form factor. The emission path uses a collection NA of 0.45 and 1.6× magnification, compared with 0.3 NA and 3× magnification in the UCLA Miniscope v4 optical path. Excitation light from a blue LED (LXZ1-PB01, LUXEON Z; 470 nm), depicted in Fig. 2c, is spectrally filtered by a bandpass excitation filter (ET470/40x, Chroma), transmitted through a shortpass dichroic beamsplitter (T500spxr, Chroma), and delivered to the sample through the objective. The objective comprises a molded polycarbonate asphere (15-273, Edmund Optics) and a plano-convex lens closest to the sample (45-226, Edmund Optics).

The same objective collects the fluorescence emission depicted in Fig. 2b. The emission is reflected through 90° by the dichroic and relayed through a second molded polycarbonate asphere serving as the tube lens (14-572, Edmund Optics). The path is folded a second time by a planar mirror (PFR10-P01, Thorlabs, diced to 5 mm × 7 mm), after which the light passes through a bandpass emission filter (ET525/50m, Chroma) and onto a monochrome CMOS image sensor (onsemi, NOIP1SN0480A-STI). Folding the path twice distributes the optical elements laterally rather than vertically, reducing the overall height from 22 mm (v4 Miniscope) to 12 mm, and the molded polycarbonate aspheres reduce the optical assembly mass. The imager employs a set-screw-based manual focusing mechanism that enables adjustment of the optical stack within the baseplate. During baseplating, this mechanism positions the optical focal plane relative to the implanted GRIN lens and then mechanically stabilizes it for subsequent recordings.

Three orthogonal coils mounted on the head-mounted assembly receive wireless power. The receiver coils feed multi-input rectification and power-management circuits that supply regulated voltages to the image sensor, microcontroller, excitation LED, and optical communication frontend. Each receiver coil weighs approximately 200 mg. A lithium-polymer battery is connected as an onboard power buffer to accommodate transient current fluctuations and temporary reductions in wireless power coupling during animal motion. Testing showed that an 11 mAh battery weighing 0.33 g was sufficient for continuous operation, yielding a total head-mounted mass of 4.12 g. A 50 mAh battery weighing 1 g was used in initial recordings and in multi-animal and complex recording sessions as a conservative choice to provide additional buffering margin against variations in head orientation and instantaneous power-transfer efficiency.

### Bench optical characterization

Bench characterization was performed to compare the optical performance of the Miniscope Zero with that of the UCLA Miniscope v4 and to quantify the spatial resolution of the Miniscope Zero optics (Fig. 2e–h). To isolate the optical system’s contribution, both configurations used an identical image-sensor PCB, with a v4 PCB mounted on each assembly. The image sensor, sensor electronics, gain, and exposure were matched across all comparisons. The miniscope was mounted on a Thorlabs micromanipulator to set and adjust the working distance relative to the imaging target. All images were acquired as raw 8-bit miniscope output and analyzed in ImageJ.

#### Emission collection

To compare emission collection, we imaged a microLED display panel (JBD, 401001B01) as an emission-only target, with no excitation light path engaged. Acquisition settings were identical across systems. For each system, a stream of frames was recorded, and a temporal average projection was computed from 200 frames. Histogram counts were converted to probability distributions, and mean per-pixel intensity within the region of interest was computed in 8-bit arbitrary units. The brightness ratio was defined as the ratio of Miniscope Zero to Miniscope v4 mean ROI intensity (Fig. 2e).

#### Spatial resolution

The spatial resolution of the Miniscope Zero was characterized by imaging a Thorlabs negative USAF 1951 resolution test target. Resolution was assessed from line-intensity profiles drawn across both the horizontal and vertical bar triplets of each element. An element was scored as resolved when the three bars produced three distinct intensity maxima separated by two local minima in the normalized profile. The smallest fully resolved element was Group 7, Element 3, corresponding to a bar width of 3.1 µm and a spatial frequency of 161.3 lp mm⁻¹; the corresponding line-pair period of 6.2 µm is reported as the resolution limit (Fig. 2f,g).

#### Point-spread function

The lateral point-spread function was measured by imaging 200 nm fluorescent microspheres (TetraSpeck, Thermo Fisher Scientific) dried onto a coverslip and imaged through the objective and emission path. Isolated beads with no neighboring bead within 23 µm (≈4× the measured full width at half maximum (FWHM)) and peak intensities below 50% of the sensor full scale were selected. Each bead was fitted with a two-dimensional Gaussian integrated over the pixel area. Across *n* = 10 beads, FWHM was 6.05 ± 0.54 µm (mean ± s.d.; FWHM was 6.09 ± 0.58 µm along the *x* axis and 6.01 ± 0.52 µm along the *y* axis).

#### Excitation profile

Spatial uniformity of the excitation field was characterized by replacing the sample with a bare Sony IMX335 monochrome image sensor (2.0 µm pixel pitch, monochrome), positioned at the nominal working distance of the excitation path; no fluorescent target or sample was used. A temporal average projection was computed from multiple frames to suppress per-frame noise. The FWHM of the illuminated field was measured along horizontal and vertical cross-sections through the intensity centroid as the outer-to-outer span of contiguous pixels at or above half-maximum, converted to physical units using the pixel pitch. Illumination uniformity was quantified as the coefficient of variation (CV = standard deviation/mean) of the background-subtracted intensity (Extended Data Fig. 1b,c).

### Simulated optical characterization

The Miniscope Zero excitation and emission optical paths were designed and evaluated using sequential ray tracing in Zemax OpticStudio (Ansys; Zemax Optic Studio 19.4 SP2). The emission path was modeled at a single design wavelength of 525 nm, within the GCaMP emission range. The emission path was evaluated at the field center and at two positions 0.5 mm from the optical axis to assess off-axis performance beyond the nominal imaging field. A physical aperture stop with a 2.7 mm semi-diameter was located in the collimated space between the dichroic and the tube lens, defining the system aperture (Fig. 2b) and yielding an object-space numerical aperture of 0.45. Lens spacings (axial air gaps between elements) were treated as variables and optimized using a damped-least-squares local optimizer to minimize a merit function that targets RMS spot size across the three field points. The sample medium was modeled as a thin seawater layer (refractive index ≈ 1.34) behind the imaging window, as a refractive-index proxy for aqueous brain tissue. Optical performance was assessed from the simulated FFT modulation transfer function and the geometric spot diagram across the field (Fig. 2i–l).

### Wireless power cavity design, construction and operation

Wireless power was delivered using a QSCR transmitter designed to generate a volumetric magnetic field over the behavioral arena (Fig. 3a,b). The cavity was designed to operate at 6.63 MHz and to couple efficiently to the three orthogonal receiver coils on the head-mounted Miniscope Zero. The cavity geometry and resonant behavior were designed in COMSOL Multiphysics using the RF Module. Simulations evaluated the magnetic field distribution and guided the design for wide-area power delivery within the experimental volume (Fig. 3b). The cavity was fabricated from A1100 aluminum using standard sheet-metal cutting and bending processes. A1100 aluminum was selected for its high electrical conductivity and compatibility with large-area fabrication. Individual panels were bolted together to maintain low-loss electrical connections across seams and reduce resistive losses during resonant operation. Lumped capacitors were mounted at defined locations to tune the resonant frequency to 6.63 MHz. An RF power amplifier based on the EPC9065 zero-voltage-switching (ZVS) class-D amplifier design (Efficient Power Conversion) drove the cavity through an inductively coupled drive loop (Extended Data Fig. 3a,b, Extended Data Fig. 7a).

During experiments, the cavity surrounded the behavioral arena and was driven continuously. Input power was adjusted to provide sufficient power at the head-mounted receiver while maintaining stable operation of the Miniscope Zero electronics. An approximate battery voltage of 3.5 V was used as an operational threshold during experiments, providing margin above the 3.3 V regulated supply rails. To evaluate wireless-power stability under in vivo operating conditions, two mice carrying Miniscope Zero devices were placed simultaneously in a 300 mm × 300 mm open-field arena and allowed to move freely. Battery voltage from each device was monitored and transmitted over the optical link. The resulting voltage trajectories were used to assess maintenance of the onboard battery buffers during simultaneous two-animal operation relative to battery-only operation under comparable payload constraints (Fig. 3f). Battery voltage was decoded from raw analog-to-digital converter values and smoothed for visualization in both this experiment and the 9 h single-animal capability test shown in Extended Data Fig. 6.

### Wireless power system evaluation

Wireless power-transfer efficiency was estimated from S-parameters measured using a vector network analyzer (VNA). One VNA port was connected to the QSCR drive coil, and the second port was connected to the head-mounted receiver coil. Measurements were performed at 6.63 MHz while the receiver was positioned at defined locations and heights within the arena.

The measured S-parameters were converted to the maximum achievable transfer efficiency using the maximum transducer-gain expression developed in prior work^31^. This calculation estimates efficiency under the receiver load condition that maximizes power transfer. To attribute transfer losses, the same measured S-parameters were fitted to a three-resonator coupled-circuit model representing the drive coil, QSCR cavity, and receiver coil. At the frequency and complex load that maximized transfer efficiency, the model currents were solved, and the resistive loss in each modeled component was expressed as a fraction of the total resonator loss, excluding power delivered to the load. The receiver matching network was tuned using series and parallel capacitors to approximate this condition. Measurements were repeated across receiver positions and heights to generate the position and height-dependent efficiency maps shown in Fig. 3e. Available receiver power was measured separately by driving the cavity and recording the DC power delivered to a 100 Ω resistor at the receiver output, with the receiver placed 50 mm above the arena floor at four positions spanning from the center to near a corner.

Because the effective receiver load varies with device operation and battery charging, input and output power were also estimated over a range of load impedances using the measured S-parameters and a voltage-source model representative of the class-D inverter that drives the QSCR transmitter. Receiver-matching capacitors were selected to provide efficient power transfer near an effective load resistance of approximately 100 Ω at representative recording positions, while maintaining usable output power across nearby variations in resistive and reactive loads.

Because tissue heating is a principal concern for radiofrequency exposure at 6.63 MHz, whole-body-average specific absorption rate (SAR) was simulated using the RF Module in COMSOL Multiphysics. The mouse was approximated as a homogeneous cylindrical phantom^35^ with a radius of 12.5 mm and a height of 60 mm, corresponding to a mass of approximately 29.5 g at a density of 1,000 kg m⁻³. Following simplified wireless-power dosimetry models, the phantom was assigned dielectric properties equal to two-thirds of the skeletal-muscle values at 6.63 MHz obtained from the IT’IS Tissue Properties Database using the Gabriel model^36–38^. This yielded a relative permittivity of 159 and an electrical conductivity of 0.401 S m⁻¹. Relative permeability was set to 1. Whole-body-average SAR was calculated from the volume-averaged electromagnetic power dissipation and normalized to a transmitter input power of 5 W. SAR was evaluated under an unloaded-receiver condition, which was treated as a high-field exposure condition at fixed transmitter input power^25^. ICNIRP occupational and general-public human-exposure limits were included for reference and were not interpreted as mouse-specific safety thresholds^39^.

### Optical data I/O modules

The Miniscope Zero communication system comprises a high-speed optical uplink for imaging data and metadata and a low-speed infrared downlink for device commands (Fig. 3a,c,d; Extended Data Fig. 7b). The optical uplink supports 8 Mbps transmission, sufficient for 8-bit imaging at 200 × 200 pixels and 20 frames per second, plus metadata for frame reconstruction and device-state monitoring. The infrared downlink supports 2 kbps transmission and uses firmware-defined device identifiers to address commands to individual Miniscopes.

For uplink transmission, imaging data and metadata were Manchester-encoded in hardware before being driven to the optical emitter. Manchester encoding embeds the clock within the transmitted waveform, permitting data recovery without a separate clock channel. The encoded waveform was generated using an XOR logic IC (Nexperia, 74LV1T86GXH) and applied to a compact optical modulation frontend. The optical emitter comprised a high-speed surface-mount red or near-infrared LED driven by a GaN FET-based modulator (EPC, EPC2038). A red emitter (Würth Elektronik, 150060RS75003) was used for single-animal recordings. For multi-animal operation, the same red emitter and a near-infrared emitter (QT Brightek, QBLP601-2IR4) provided wavelength-separated channels.

Optical uplink signals were detected by silicon photomultiplier (SiPM)-based receiver modules (Hamamatsu, C15522-3010SA) positioned above or around the behavioral environment. To reduce electromagnetic interference from the wireless power field, each SiPM frontend was housed in a custom CNC-milled aluminum enclosure. The shielded SiPM output was routed to a custom high-gain amplifier and an adaptive digitizer that compensated for variations in the received optical intensity caused by animal position, head orientation, and line of sight. An FPGA-based detector (Opal Kelly, XEM7310) synchronized to the Manchester-encoded signal, performed clock recovery and data decoding, and recovered the transmitted packets. Decoded data were transferred to the acquisition computer through a USB interface.

Multiple receiver modules could be operated in parallel to provide redundant coverage of large or geometrically complex environments. Frame reconstruction and post hoc combination of receiver streams are described below.

Receiver filtering was configured according to the experimental mode. For single-animal recordings, receivers used filters matched to the uplink wavelength while rejecting behavioral illumination. For multi-animal recordings, wavelength-division multiplexing was implemented using red and near-infrared uplinks, with separate optical bandpass filters to isolate the corresponding spectral bands.

The infrared downlink used amplitude shift keying modulation on a 38 kHz carrier. Downlink commands adjusted sensor gain, excitation LED intensity, imaging region of interest, and other device parameters during ongoing experiments. The downlink also supported remote hardware reboot.

### Data stream packet structure

Optical uplink data were transmitted as a sequence of image-buffer packets. Each packet consisted of a fixed preamble, an 11-word header, and an image payload. A short sequence of dummy 32-bit words preceded the preamble to allow the Manchester decoder to stabilize its clock recovery. Header entries were 32-bit words encoding packet and device metadata, including frame index, global packet index, packet index within the frame, device timestamp, pixel count, raw battery voltage, and raw input voltage. During host-side parsing, battery and input voltages were converted to physical units using constants defined in the device configuration.

Each image-buffer payload contained 5,032 8-bit pixel values. A 200 × 200 pixel frame was transmitted as eight image-buffer packets. Our data acquisition software used the frame index, global packet index, within-frame packet index, payload length, and packet-ordering information to associate packets with the correct frame, device, and acquisition stream, assemble complete frames, and detect missing or duplicated packets. Host-side acquisition added runtime metadata including packet reception time. This metadata was used for post hoc alignment of the exported receiver video streams and for acquisition diagnostics.

### Packet error rate evaluation

A dedicated Miniscope Zero firmware mode was used to evaluate packet error rate (PER). In this mode, a pseudorandom binary sequence replaced the imaging data within each image-buffer payload. The test sequence was generated using PRBS-15 with polynomial x¹⁵ + x¹⁴ + 1. Each transmitted buffer contained 5,032 8-bit payload values, corresponding to 40,256 payload bits. The PRBS sequence was reseeded at each buffer boundary using seed = (buffer count mod 2¹⁵) ∨ 1, where buffer count was the monotonically increasing buffer index. The ∨ 1 operation prevented the all-zero linear feedback shift register state. The host computer regenerated the expected sequence from the transmitted buffer counter and compared it with the received payload.

A packet was classified as failed if any received payload bit differed from the expected sequence. The reported PER therefore provides a conservative measure of optical link reliability at the image-buffer level because a single bit error within the 40,256-bit payload caused the complete packet to be classified as failed. For each measurement condition, one trial comprised 2¹⁵ − 1 transmitted buffers, corresponding to one complete PRBS-15 phase cycle. PER was calculated as the number of failed packets divided by the total number of evaluated packets. Five trials were performed for each condition, and the reported PER was averaged across the five trials.

### Frame reconstruction and quality control

The data acquisition software associated image buffer packets with the corresponding frame, device, and acquisition stream using the transmitted packet metadata and reconstructed a separate video stream for each optical receiver. During acquisition, the real-time viewer applied FFT-based low-pass filtering to suppress stripe-like image noise for online visualization. This filtering affected only the displayed images and was not applied to the exported data used for offline analysis. Calcium traces shown in the figures were low-pass filtered for display (zero-phase Butterworth, 0.9 Hz, second order); spatial-coding analyses used the unfiltered traces. Each receiver stream was exported as an AVI file, along with a CSV metadata file containing the headstage-generated frame counter and packet reception times, and runtime logs with host-side timestamps and reconstructed frame indices.

For single-animal recordings acquired by two optical receivers, the exported receiver videos and metadata were combined offline. Candidate frames were aligned using the shared headstage-generated frame counter and packet reception times. When both receivers provided candidate reconstructions for the same frame, a quality score based on metadata consistency, edge artifacts, and contiguous black regions due to padding of unrecovered pixels was used to select the reconstruction with fewer missing or corrupted pixels.

Frames in the single-receiver and offline-combined videos were then screened for artifacts caused by optical-link or packet-recovery errors. These appeared as partially or fully corrupted frames, including corruption distributed across the image, and as black or incomplete frames resulting from a failure to detect the packet preamble or to recover the image payload. Frames were scored automatically in MIO from the reconstructed images and their metadata, and the affected frames were rejected before downstream analysis. Where visible corruption survived the automated screen, the recording was inspected frame by frame, and the remaining affected frames were rejected manually; this manual step was performed independently for the analysis and for the Supplementary Video 1 material. For the illustrative timelines in Extended Data Fig. 4e, missing-frame events were identified from gaps exceeding 75 ms between the timestamps of frames retained after timestamp validation, in each receiver stream and in the offline-combined stream.

### Frame transmission loss evaluation during wireless recordings

Frame transmission loss quantifies packet-level delivery only. It was quantified from the per-packet metadata logs written by each receiver during acquisition; image data were not used. The headstage transmitted each frame as eight packets, and each packet header contained the headstage frame number identifying the corresponding frame. Packets with corrupted or otherwise unreadable frame numbers were discarded. For each receiver, a frame was considered received only when all eight packets could be assigned to that frame; otherwise, it was considered lost. Frames for which all eight packets were received were not classified as lost even when the reconstructed image contained payload corruption, because isolated incorrect pixel values could not be distinguished reliably from normal image variation. Frames with visually apparent or large-scale corruption were handled separately by the frame quality-control procedure described above and were not included in the frame transmission-loss metric.

For single-animal recordings acquired by two receivers, a frame was classified as lost from the combined stream only when it failed the complete-packet criterion on both receivers. The analyzed frame range was restricted to the headstage frame-counter span shared by both receivers. For two-animal recordings, each receiver captured the wavelength channel corresponding to a different animal and was analyzed independently using its own frame-counter span.

For all recordings, the number of expected frames was calculated from the inclusive headstage frame-counter span within the analyzed interval, thereby including frames for which no packets were received. Frame transmission loss rate was calculated as the number of lost frames divided by the number of expected frames. The first 30 min of each recording were analyzed for the environment and for two-animal comparisons, as shown in Extended Data Fig. 5a,b. The full 3 h complex-environment recording and the full 9 h recording were analyzed for Fig. 6c and Extended Data Fig. 6, respectively. For the time-resolved visualizations in Fig. 6c and Extended Data Fig. 6, frame transmission loss rates were calculated in 0.5 min bins and smoothed with a Gaussian kernel with a 5 min full width at half maximum.

### GRIN lens imaging in freely behaving mice

For open-field place-cell recordings, mice carrying Miniscope Zero were recorded in a 500 × 500 mm square open-field arena made of white polypropylene. A large black visual cue was placed on each of the four walls to provide stable spatial-orientation cues. The arena was housed within the QSCR, which wirelessly powered the Miniscope Zero. Animals were filmed from above using an overhead ELP 8 MP USB wide-angle camera under blue-light illumination. Two silicon photomultiplier data-acquisition detectors were mounted above the arena, with each detector covering approximately half of the floor. The optical data receivers and behavioral camera were synchronized using an NTP server connected over Ethernet. To spectrally separate behavioral illumination from the Miniscope optical data signal, Miniscope Zero transmitted data optically over a red channel, with red filters installed on the detectors and blue behavioral illumination used.

Open-field recording sessions lasted 30 min, and small droplets of rodent yogurt were scattered across the arena floor to encourage exploration. Animals were habituated to head fixation, which was required for mounting the Miniscope, and to carrying the head-mounted microscope. Mice were first habituated to the box over five daily 10-min sessions. Place-cell recordings were then performed across five daily sessions, with each animal recorded once per day. One additional 60-min open-field session was acquired for the example shown in Fig. 4.

For enclosed-maze recordings, mice were imaged as they navigated an enclosed maze with rooms and overlapping arms (Fig. 5a,b). The maze geometry was designed to create a mechanically constrained behavioral environment in which tethered head-mounted microscopes would be prone to cable interference. Mice were recorded while freely traversing the maze under wireless power and optical data transmission. Because the animal was not always visible through the maze structure, position was tracked from the Miniscope data LED rather than from body-part pose.

### Complex-environment and two-mouse multilevel recordings

For the complex-environment demonstration, a mouse carrying Miniscope Zero was recorded in an arena containing a multilevel structure, stairs, a running wheel, and open floor space (Fig. 6a). The arena was positioned within the QSCR cavity, and the microscope was operated under wireless power with optical data transmission throughout the session. Because the three-dimensional structures intermittently occluded the animal, behavioral position was tracked from the Miniscope data LED rather than from body-part pose. Areas corresponding to the multilevel structure, wheel, stairs, and floor were manually defined from the behavioral video, and the LED trajectory was used to assign the animal to behavioral zones over time.

For the two-mouse multilevel demonstration, two animals carrying Miniscope Zero devices were recorded simultaneously in a multilevel arena using wavelength-separated optical data channels (Fig. 6i). Animals were identified by separate red and near-infrared data LEDs and tracked independently. A multi-animal DLCRNet model with a ResNet-50 backbone, output stride 16, five multi-fusion stages, and a bottom-up part-affinity-field architecture was trained for 800 epochs to detect the two LED markers. Identities were maintained using DeepLabCut’s identity head and a transformer re-identification model.

The two-mouse multilevel pipeline was evaluated for method validation rather than for statistical comparison, and no inferential statistical tests were applied. For each animal, per-frame level assignment was used to quantify inter-level transitions. Detection accuracy was validated against manually annotated transitions verified by frame-by-frame inspection of the video. To suppress spurious level changes caused by tracking jitter near level boundaries, per-frame level labels were smoothed using a 2-s sliding-window majority vote followed by a 4-s minimum-dwell filter; both thresholds were calibrated against the ground-truth annotations.

### Miniscope behavior comparison

Behavioral effects of carrying head-mounted devices, which differed in both mass and tethering, were assessed in a 30 × 30 cm square open-field arena made of white polypropylene (Fig. 4e–g). Animals were filmed from above using an ELP 8 MP USB wide-angle camera under uniform diffuse white light. To encourage exploration, small hand-cut fragments of rodent yogurt drops were scattered randomly across the arena floor before each 15-min recording. To isolate the mechanical effects of mass and tethering, these sessions were run with the QSCR transmitter deliberately unpowered, so that both scope conditions were compared under identical environmental conditions.

Two head-mounted microscope systems were compared. Calcium imaging was not performed in these behavioral comparison experiments, but the flexible coaxial cable was attached during Miniscope v4 recordings to replicate the mechanical tethering used in conventional imaging. Miniscope Zero was tested with a dummy headstage, comprising the Miniscope body, wireless power receiver coils, and an 11 mAh battery, for a total head-mounted mass of 4.12 g. The Miniscope v4, likewise non-operating, weighed 2.8 g without its tether. Thus, the comparison contrasted a lighter but tethered system with a slightly heavier but cable-free system. A single Miniscope Zero baseplate was used for each animal, and a custom 3D-printed adapter allowed Miniscope v4 to mount onto the same baseplate.

After four sessions of arena habituation, mice were recorded across three no-scope sessions, whose median defined the per-animal control baseline. At this stage, animals had not been exposed to wearing either microscope. Animals were then habituated over six sessions to carrying Miniscope Zero and Miniscope v4. This was followed by four comparison sessions in which each mouse was tested with both miniscopes. Miniscope assignment was counterbalanced across animals and recording days. One mouse died during the experiment and was excluded from all analyses, leaving nine animals.

### Behavioral video correction and position tracking

Behavioral video frames were corrected for radial barrel distortion using OpenCV and then perspective-rectified to a square using four manually selected arena corners. Distortion coefficients and corner coordinates were tuned interactively using a calibration frame and adjusted for each recording day or camera configuration.

For open-field place-cell, two-animal multilevel, and behavioral-comparison recordings, animal position was tracked with DeepLabCut version 3.0.0rc1 using the PyTorch engine. For open-field place-cell recordings, the position of a single mouse was tracked from a red data-transmitting LED mounted on Miniscope Zero. A single-animal model with a ResNet-50 backbone and a heatmap prediction head with location refinement was trained for 200 epochs to detect the LED. For two-animal recordings in the multilevel arena, the animals were distinguished by the wavelengths of their data-transmitting LEDs. The red and near-infrared LEDs were tracked using a multi-animal DLCRNet model.

To compare Miniscope Zero and Miniscope v4 head-mounted loads (Fig. 4e–g), animals carried non-functional dummy miniscopes without a data-transmitting LED, so their positions were tracked from body landmarks. A top-down model comprising an animal detector followed by a pose estimator with an HRNet-W32 backbone was used to track the body center, nose, and tail base. Body-center coordinates with likelihood below 0.6 were replaced by the midpoint of the nose and tail base when both landmarks were confidently detected; otherwise, the body center was estimated from the tail-base position using the median body-center-to-tail-base displacement.

For enclosed-maze and complex three-dimensional environment recordings, conventional body-part pose estimation was not applicable because the mouse was not continuously visible throughout the arena. The Miniscope data LED was therefore tracked using a custom single-keypoint convolutional neural network with an ImageNet-pretrained ResNet-18 backbone and a transposed-convolution decoder. RGB frames were resized to 224 × 224 pixels, and the network predicted a 56 × 56 Gaussian heatmap. LED position was estimated from a weighted combination of the heatmap peak and its intensity-weighted centroid. The network was trained using a combined heatmap mean-squared-error and peak-coordinate loss for up to 1,000 epochs, with early stopping, a 20% validation split, and a learning rate of 5 × 10^−4^.

### Trajectory preprocessing

Animal position was processed using a standard geometric preprocessing pipeline comprising marker cleaning, parallax correction, clipping to the arena boundaries, conversion to physical units when applicable, and speed computation.

For open-field place-cell recordings, raw trajectories were cleaned with a Hampel filter. At each frame, position was compared with the centered rolling median over a 7-frame window, and samples deviating by more than 3 × 1.4826 × MAD were flagged as outliers and linearly interpolated. No likelihood threshold was applied.

For enclosed-maze recordings, raw x, y tracks were first denoised using a Hampel filter based on a rolling spatial median and MAD-based outlier rejection, with flagged jumps replaced by linear interpolation. Rooms were defined as polygons and arms as polylines. Each frame was assigned to the highest-probability zone under a soft-boundary model that combined signed distance to a room polygon, or perpendicular distance to an arm centerline, with distance-decay weighting. Transitions were constrained by a precomputed adjacency graph linking each room to the arms it opened onto, so room and arm transitions followed the physical maze topology. When assigned to an arm, position was projected onto the arm centerline to yield a normalized arc-length coordinate between 0 and 1.

For the 9 h in vivo capability test shown in Extended Data Fig. 6, locomotion speed was calculated from tracking data acquired at approximately 20 frames per second using a 0.25 s displacement window, then downsampled to 1 Hz and median-filtered over 5 samples.

Because the tracked point was above the arena floor while videos were acquired from overhead, points away from the image center were projected radially outward in proportion to marker height. Each position was therefore scaled radially toward the arena center by a factor of (H − h)/H, where H is the camera height above the floor, and h is the height of the tracked point on the animal. Arena bounds were rescaled by the same factor before conversion to real-world units. The corrected trajectory was clipped to arena bounds. For open-field place-cell recordings, data were converted to physical units using the known arena dimensions. Movement speed was computed from corrected positions over a centered 0.25-s window. Frames below a running-speed threshold of 25 mm/s were excluded from place-cell analyses.

### Neural data preprocessing

Videos retained after frame reconstruction and quality control were preprocessed offline before cell detection. Horizontal readout stripes were first removed in ImageJ using a directional FFT filter that suppresses stripe noise. The destriped videos were then processed with a custom Python pipeline that removed frame-wide intensity offsets by per-frame median correction, subtracted a per-pixel minimum-intensity projection to remove static background fluorescence and fixed-pattern noise, applied Gaussian smoothing and morphological top-hat filtering to suppress residual noise and time-varying background, and scaled the result with a constant gain for a given animal and paradigm. The resulting videos were processed with MiniAn, including motion correction, before cell detection.

Neuronal regions of interest were identified from the processed videos using a MiniAn-based cell detection pipeline. MiniAn’s temporal update solves for the denoised calcium trace and the deconvolved activity jointly, so denoising cannot be separated from deconvolution. This step was bypassed: per-ROI traces were taken without autoregressive denoising. Calcium event activity was then estimated using OASIS deconvolution before spatial coding analyses.

### Open-field spatial tuning analysis

Spatial coding in the open field was quantified from frames in which running speed exceeded 25 mm s⁻¹. The arena was divided into a 50 × 50 grid, corresponding to 1 × 1 cm spatial bins. For each recording, an occupancy map was computed as the time spent in each bin. For each unit, an event map was computed as the amplitude-weighted sum of deconvolved calcium events in each bin. The event and occupancy maps were smoothed independently using a two-dimensional Gaussian kernel with σ = 3 bins, and the activity-rate map was calculated by dividing the smoothed event map by the smoothed occupancy map. Bins with values below 0.05 s in the smoothed occupancy map were excluded. This threshold was based on minimum occupancy criteria used in related studies to exclude sparsely sampled bins^11,40^. Spatial coverage for each recording was calculated as the percentage of spatial bins the animal entered. This measure of behavioral sampling is distinct from the occupancy criterion above, which is applied to the smoothed occupancy map.

### Enclosed-maze spatial tuning analysis

Spatial coding in the enclosed maze was quantified from arm-traversal frames in which running speed exceeded 25 mm s⁻¹; periods spent in rooms were excluded. Each frame was assigned a within-arm position and traversal direction. Within each arm, position was divided into an integer number of equal bins, with the bin count selected to produce a bin width as close as possible to 10 mm. The bin sequences from each arm were concatenated in a fixed order to form a linearized spatial axis in millimeters. Because each arm was binned independently, arm boundaries coincided with bin edges and no bin spanned a junction. Forward and reverse traversals were treated as separate segments.

For each recording, an occupancy map was computed as the time spent in each linearized bin. For each unit, an event map was computed as the amplitude-weighted sum of deconvolved calcium events in each bin. The event and occupancy maps were smoothed independently using a one-dimensional Gaussian kernel with σ = 2 bins, and the activity-rate map was calculated by dividing the smoothed event map by the smoothed occupancy map. Smoothing was performed separately within each segment to prevent activity from being blurred across arm boundaries. Bins with values below 0.05 s in the smoothed occupancy map were excluded using the same occupancy criterion as in the open-field analysis. Spatial coverage for each recording was calculated as the percentage of linearized arm-direction bins the animal entered.

### Spatial information, stability and place cell classification

For both the open-field and enclosed-maze analyses, spatial tuning was summarized using Skaggs spatial information. Statistical significance was assessed using a circular-shift permutation test. Calcium events were circularly shifted relative to the trajectory by at least 20 s, and spatial information was recomputed for 1,000 shuffles. Permutation *P* values were calculated as the number of shuffled values equal to or greater than the observed value, plus one, divided by the number of shuffles plus one. Spatial information was considered significant at *P* < 0.05.

Spatial stability was assessed from correlations between activity-rate maps generated using two complementary temporal partitions: a first-half versus second-half split and an interleaved split in which the recording was divided into ten consecutive blocks assigned alternately to two groups. For each split, the Pearson correlation between the two maps was calculated and Fisher z-transformed. Significance was assessed using 1,000 circular-shift permutations with a minimum shift of 20 s and a threshold of *P* < 0.05. A unit was classified as a place cell only if it exhibited significant spatial information and passed both spatial-stability tests.

### Behavioral-impact analysis

Three metrics were computed from the cleaned trajectory for each recording: total distance traveled, locomotion speed, and immobility time. Total distance traveled was computed as the sum of path lengths in meters. Locomotion speed was computed as the median instantaneous speed across frames in which the animal moved at least 2 cm/s, thereby excluding immobility. Immobility time was computed as total time below 2 cm/s, in minutes.

Because each mouse experienced all three conditions (no scope, Miniscope Zero, Miniscope v4), behavior was analyzed within the animal. For each metric, each animal’s sessions were collapsed to a single median per condition, yielding one value per animal per condition (n = 9 animals). Group values and box-plot center lines represent the median across animals of these per-animal medians. The three conditions were compared for each metric using a Friedman test, corresponding to a nonparametric repeated-measures ANOVA. When the omnibus test was significant at *P* < 0.05, the three pairwise comparisons were assessed using two-sided Wilcoxon signed-rank tests, with *P* values corrected across the three pairs using the Holm–Bonferroni method.

### Use of artificial intelligence tools

Generative AI tools (OpenAI GPT and Claude) were used to assist software development and refine the language and grammar of the manuscript.

## Supporting information

Representative Miniscope Zero recordings across experimental environments.

## Data availability

Experimental data generated in this study are available via UCLA Dataverse: figure source data and hardware characterization (https://doi.org/10.25346/S6VGDP7C); open-field arena recordings (https://doi.org/10.25346/S6PNI5WP); behavior tracking and histology (https://doi.org/10.25346/S6ZWUJWY); enclosed maze, multilevel arena, and complex environment recordings (https://doi.org/10.25346/S6684SQV).

## Code availability

Designs for the described Miniscope Zero hardware and firmware are available at https://github.com/miniscope/MiniscopeZero. Software is available as follows: data acquisition and preprocessing (https://github.com/miniscope/mio, https://github.com/miniscope/miniscope_preproc); spatial tuning analysis (https://github.com/miniscope/camap); LED position tracking through translucent objects (https://github.com/miniscope/FuzzyTrack).

## Acknowledgments

Illustrations used throughout the figures were adapted from drawings by Annie Park, Tom Baden, and Roberta Schellino available through SciDraw (https://doi.org/10.5281/zenodo.10940481; https://doi.org/10.5281/zenodo.3926533; https://doi.org/10.5281/zenodo.10390020) under the Creative Commons Attribution 4.0 International license.

## Author information

These authors contributed equally: Takuya Sasatani, Marcel Brosch.

### Contributions

T.S., M.B., and D.A. conceived and designed the study. T.S. led wireless power development, electronics development, and embedded system integration. M.B. led optomechanical development and the behavioral-comparison study. T.S. and M.B. co-led the wireless data transfer development, analysis software development, firmware development, in vivo recordings, data analysis, and system evaluation. J.S. contributed to software development. Z.D. developed the FPGA data decoder. F.S.J. and M.S. contributed to initial in vivo testing. F.S.J. also assisted with initial prototype development and characterization. F.S.J., H.S., A.G., H.C., K.K., and B.A.M. contributed to early exploration and validation of system components. P.Z., M.S., and L.O. contributed to animal preparation, surgeries, and behavioral experiments. T.S. and M.B. wrote the manuscript with extensive input and editing from D.A. D.A., T.S., A.J.S., A.K.C., and P.G. supervised the project and provided resources. All authors reviewed and approved the final manuscript.

---

**Extended Data Figure 1.**
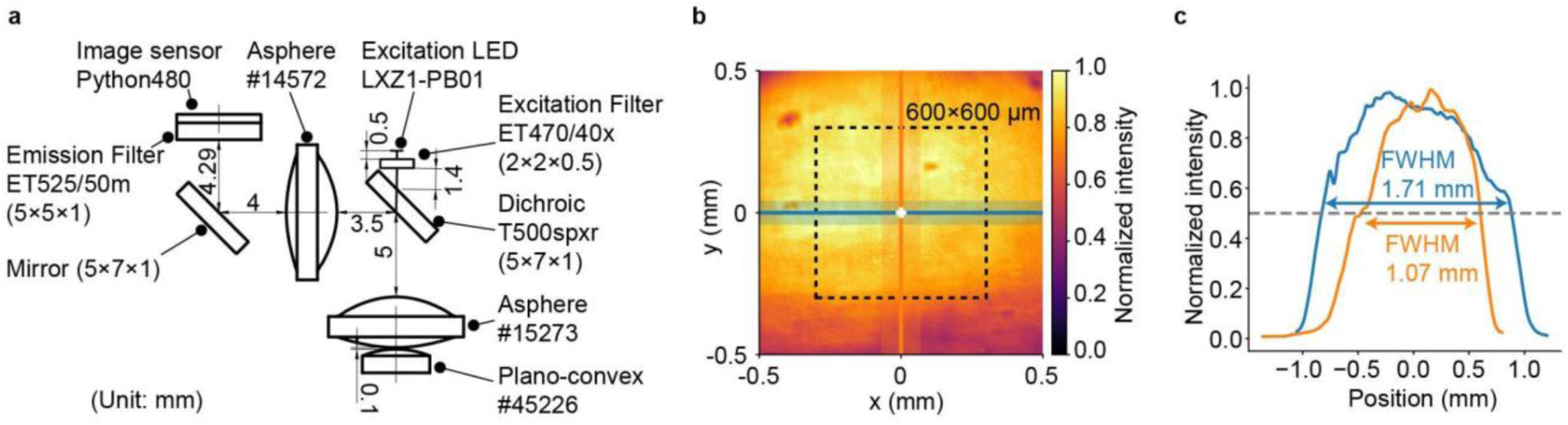
Optical layout and excitation profile of Miniscope Zero. **a,** Cross-sectional layout of the shared excitation and emission optical paths. **b,** Measured excitation-intensity distribution over a 1 mm × 1 mm analysis region at the sample plane. The dashed outline indicates the 600 μm × 600 μm imaging field, and the horizontal and vertical lines indicate the profiles shown in (c). The coefficient of variation within the imaging field was 0.07. **c,** Normalized horizontal and vertical excitation-intensity profiles through the center of the imaging field. The full widths at half maximum (FWHM) were 1.71 mm horizontally and 1.07 mm vertically.

**Extended Data Figure 2.**
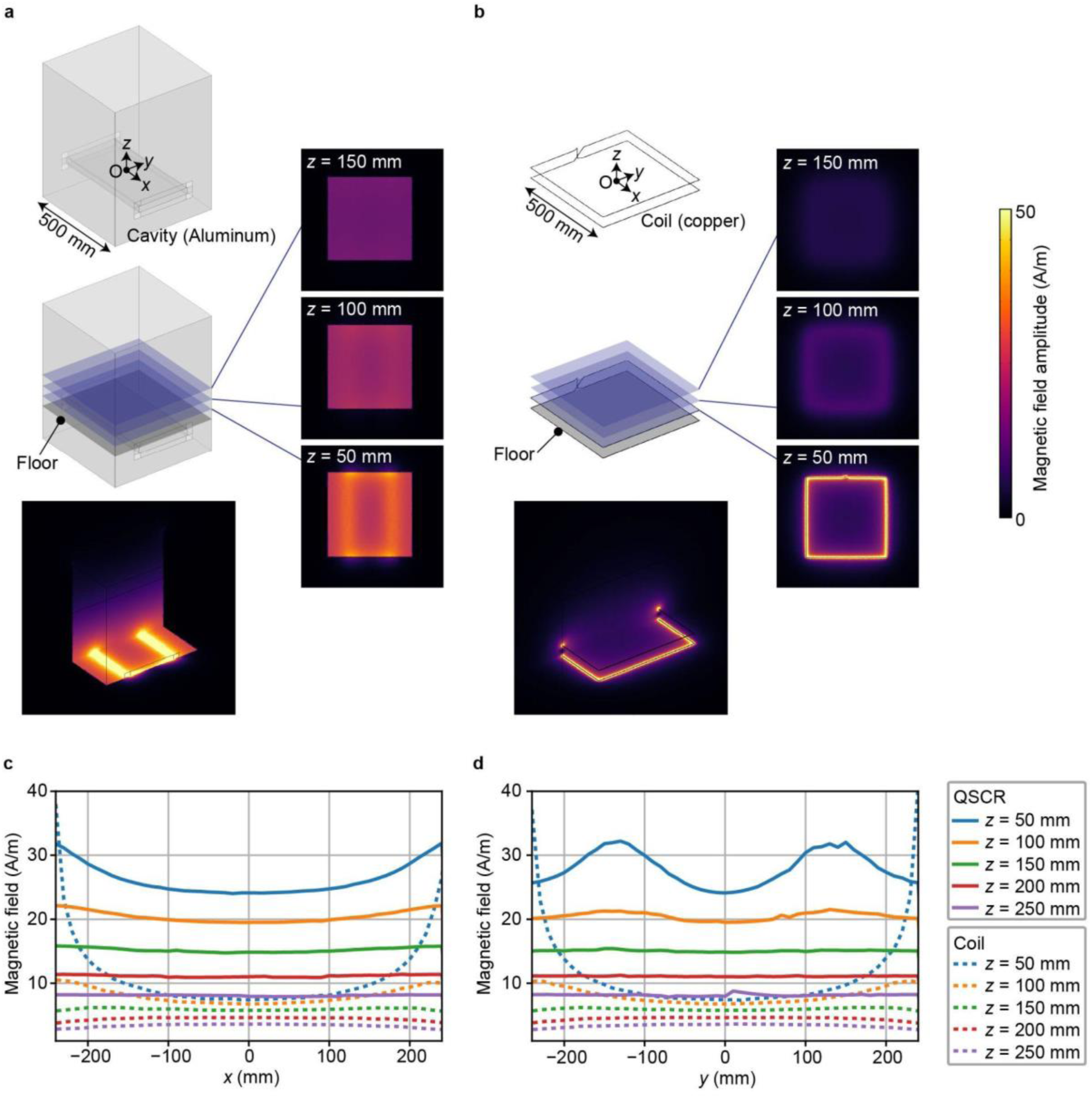
Simulated magnetic-field distribution of the QSCR and a reference coil. **a,b,** Geometries and simulated magnetic-field amplitude distributions for the QSCR cavity (**a**) and reference planar coil (**b**) at 6.63 MHz. Field maps are shown at horizontal planes 50, 100, and 150 mm above the arena floor, with matched transmitter-loss normalization. **c,d,** Magnetic-field amplitude profiles along the *x* (**c**) and *y* (**d**) axes. Solid lines indicate the QSCR cavity and dashed lines indicate the reference coil. The QSCR generates a stronger, more spatially uniform magnetic field throughout the behavioral volume.

**Extended Data Figure 3.**
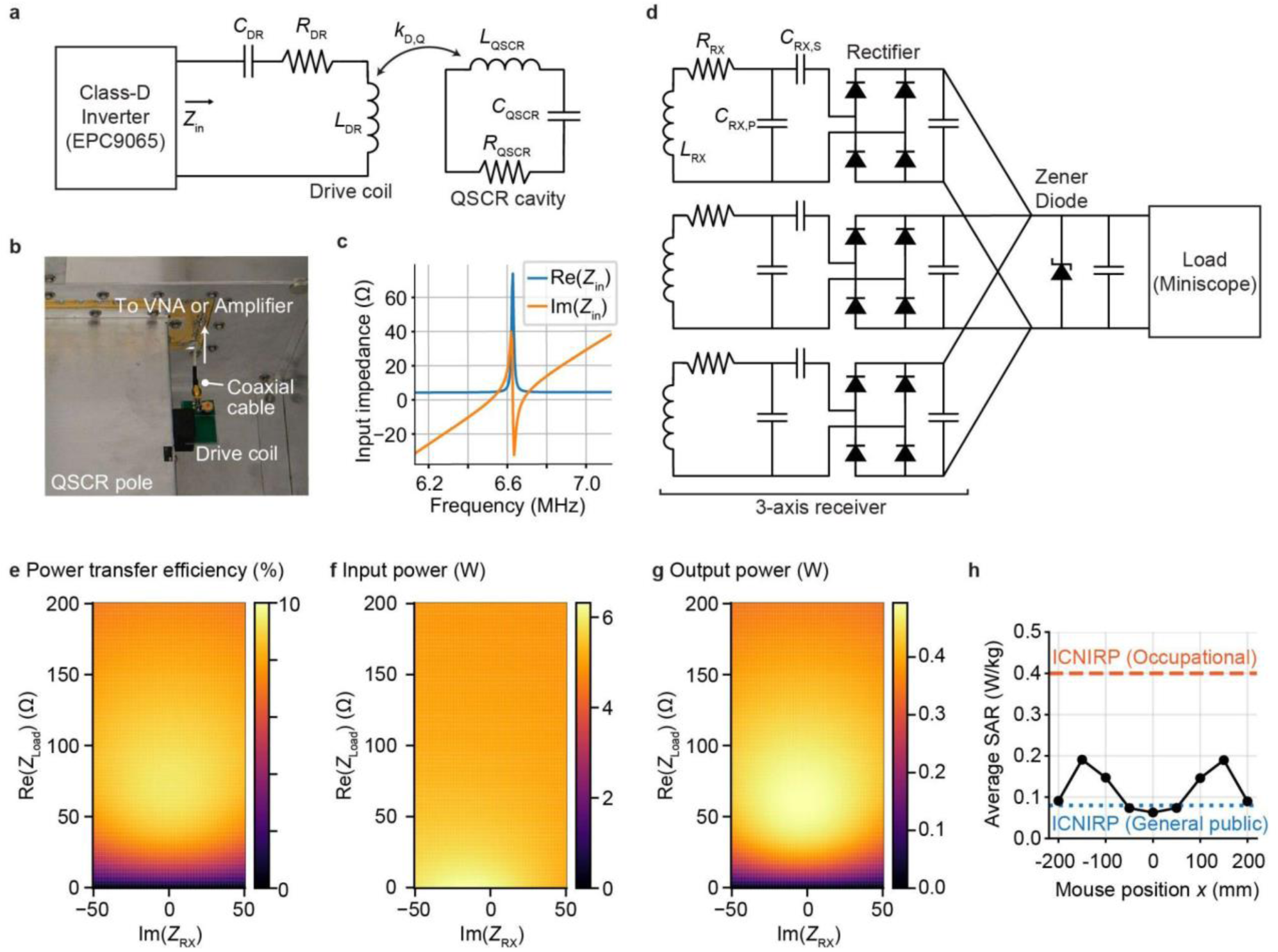
QSCR transmitter, receiver circuitry and wireless-power characterization. **a,** Equivalent circuit of the QSCR transmitter, in which a class-D inverter drives the cavity through an inductively coupled drive coil and tuning network. **b,** Drive-coil interface mounted on the QSCR pole, with a coaxial connection for amplifier drive or vector network analyzer measurements. **c,** Measured transmitter input impedance near the 6.63-MHz operating frequency. **d,** Three-axis wireless-power receiver architecture. Orthogonal receiver coils are independently tuned and rectified, and their outputs are combined and filtered before powering the Miniscope. **e–g,** Power-transfer efficiency (**e**), transmitter input power (**f**), and receiver output power (**g**) estimated from measured S-parameters across receiver-load impedances using a voltage-source model representative of the class-D inverter. **h,** Simulated whole-body-average specific absorption rate as a function of mouse position under an unloaded-receiver condition at a transmitter input power of 5 W. The mouse was represented as a homogeneous cylindrical phantom with dielectric properties equal to two-thirds of skeletal-muscle values at 6.63 MHz^35–38^. ICNIRP human-exposure limits are shown for reference^39^.

**Extended Data Figure 4.**
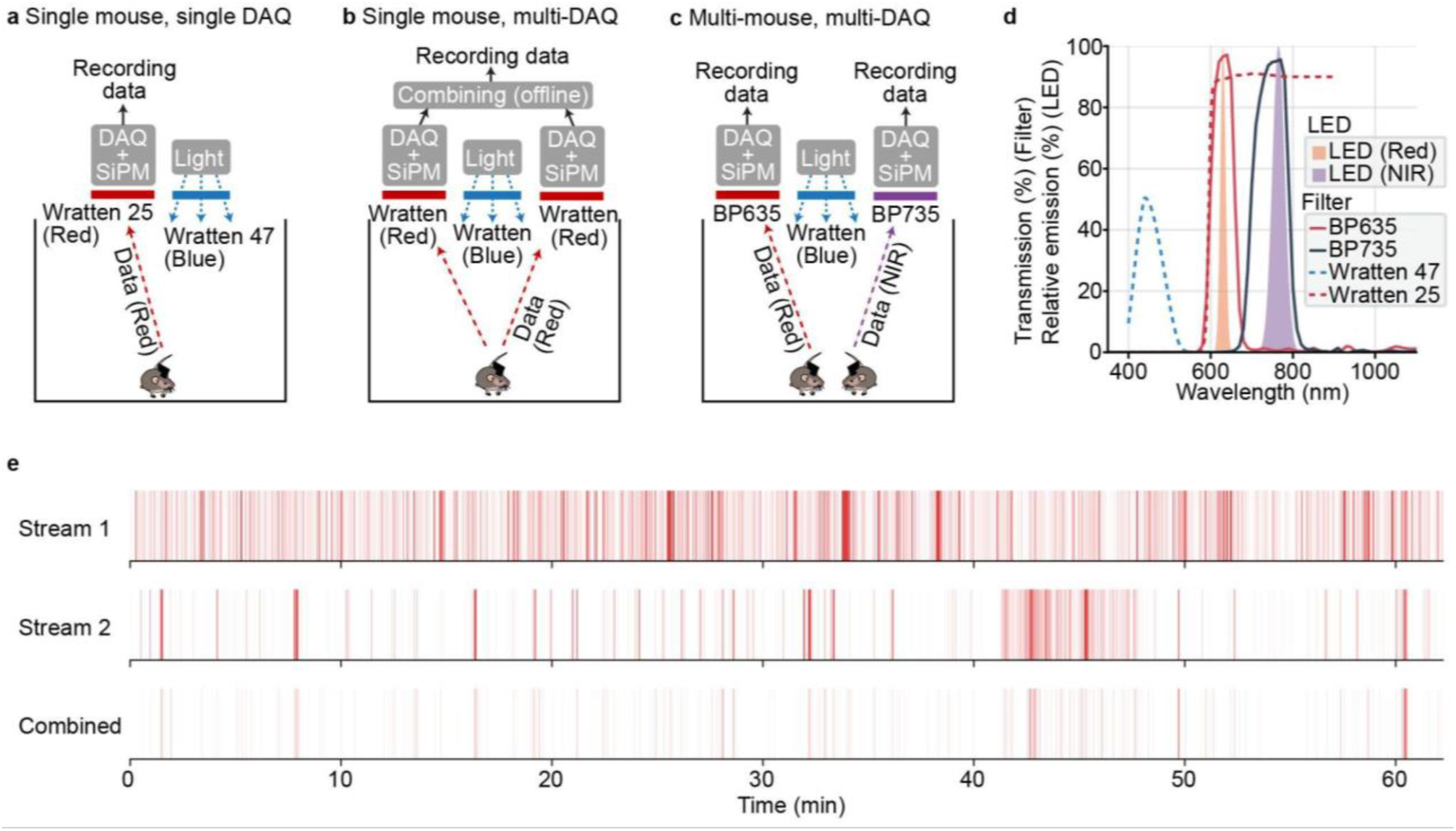
Optical acquisition configurations and dual-receiver frame combining. **a–c,** Optical receiver configurations for a single mouse with one receiver (a), a single mouse with two receivers and offline stream combining (b), and two mice using wavelength-separated red and near-infrared uplinks with separate receivers (c). **d,** Emission spectra of the red and near-infrared data LEDs and transmission spectra of the corresponding receiver filters and blue behavioral-illumination filter. **e,** Post-quality-control missing-frame events during the enclosed-maze recording shown in Fig. 5. The upper two rows show gaps in the frames retained from the individual receiver streams, and the lower row shows gaps remaining after offline stream combining and quality control. Mouse illustrations adapted from SciDraw under CC BY 4.0; full attribution is provided in the Acknowledgments.

**Extended Data Figure 5.**
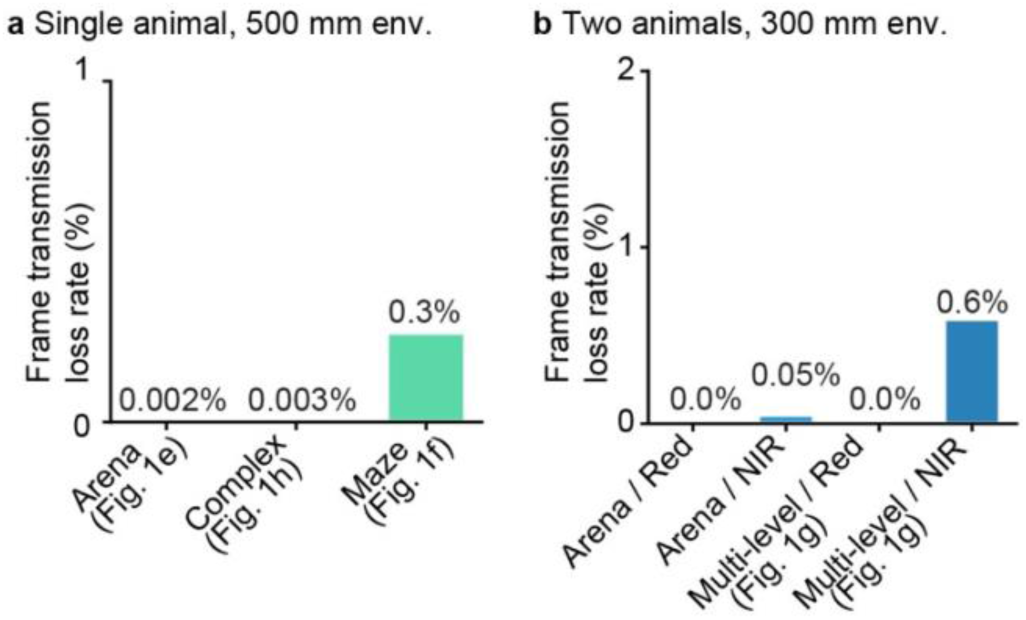
Frame transmission loss rates during single- and two-animal wireless recordings. **a,** Combined frame transmission loss rates during the first 30 min of single-animal recordings in the open-field arena, complex environment, and enclosed maze. For the combined stream, a frame was classified as lost only when neither receiver received all eight packets. **b,** Single-receiver frame transmission loss rates during the first 30 min of simultaneous two-animal recordings in the open-field and multilevel arenas, shown separately for the red and near-infrared (NIR) optical channels.

**Extended Data Figure 6.**
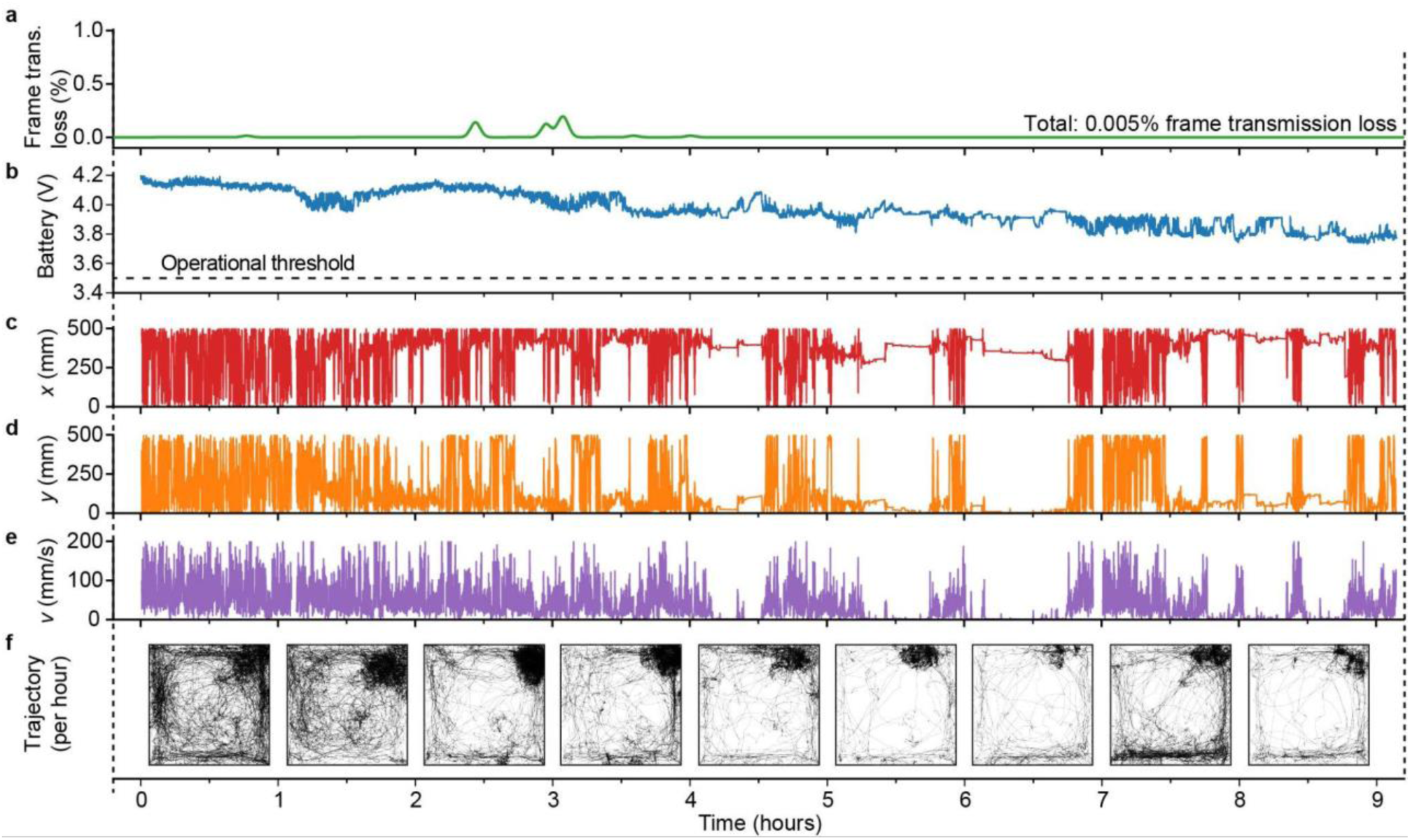
In vivo capability test of long-duration Miniscope Zero operation. A freely behaving mouse carried an operating Miniscope Zero for approximately 9 h in a 500 mm × 500 mm open arena, during which optical data packets and device telemetry were transmitted continuously. **a,** Frame transmission loss rate of the combined optical data stream, smoothed with a Gaussian kernel with a 5 min full width at half maximum; total frame transmission loss was 0.005% over the full session. **b,** Battery voltage during the recording. The dashed line indicates the 3.5 V operational threshold used during experiments. **c,d,** Tracked *x* and *y* positions of the mouse in arena coordinates. **e,** Locomotion speed. **f,** Movement trajectories shown separately for each hour of the session.

**Extended Data Figure 7.**
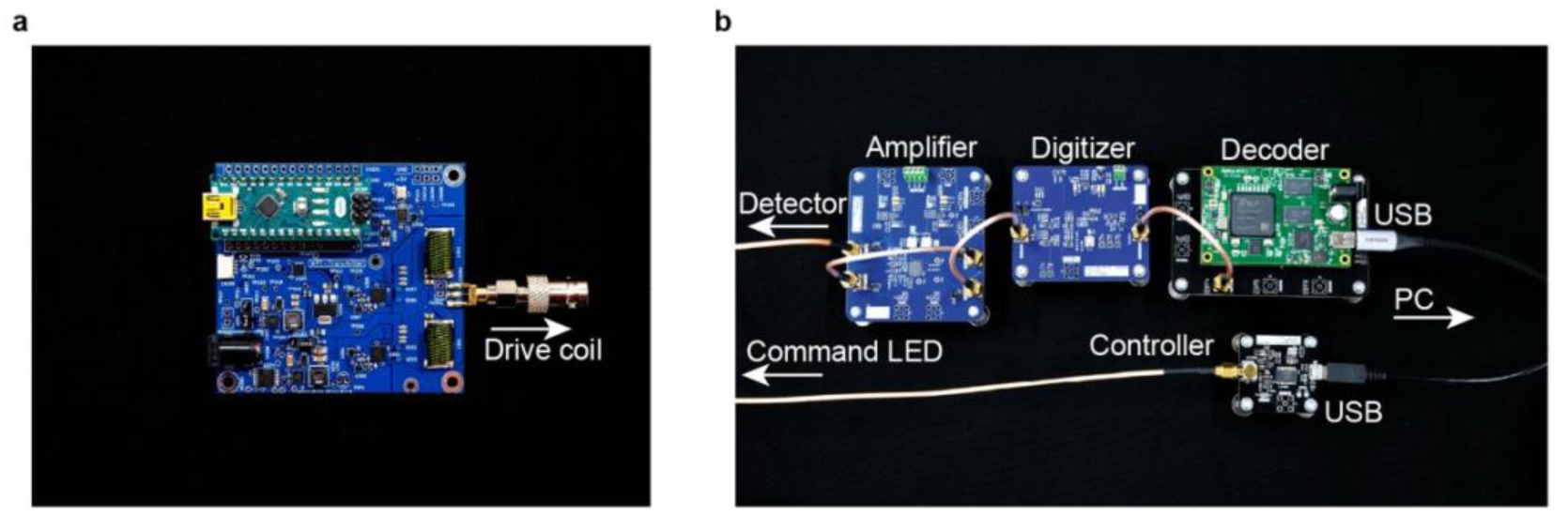
External wireless-power, data-acquisition, and command transmitter hardware. **a,** Class-D RF power amplifier used to drive the QSCR cavity through the inductively coupled drive coil. **b,** Wireless optical data-acquisition and command interface. Optical uplink signals from the detector are amplified, digitized and decoded before transfer to the acquisition PC over USB. A separate controller drives the command LED for the infrared downlink.

